# A spectrum of altered non-rapid eye movement sleep in schizophrenia

**DOI:** 10.1101/2023.12.28.573548

**Authors:** Nataliia Kozhemiako, Chenguang Jiang, Yifan Sun, Zhenglin Guo, Sinéad Chapman, Guanchen Gai, Zhe Wang, Lin Zhou, Shen Li, Robert G. Law, Lei A. Wang, Dimitrios Mylonas, Lu Shen, Michael Murphy, Shengying Qin, Wei Zhu, Zhenhe Zhou, Robert Stickgold, Hailiang Huang, Shuping Tan, Dara S. Manoach, Jun Wang, Mei-Hua Hall, Jen Q. Pan, Shaun M. Purcell

**Author notes:** corresponding authors (Jen Q. Pan,; Shaun M. Purcell,). co-first authors (Nataliia Kozhemiako, Chenguang Jiang). co-senior authors (Jun Wang, Mei-Hua Hall, Jen Q. Pan, Shaun M. Purcell).

## Abstract

**Background:** Multiple facets of sleep neurophysiology, including electroencephalography (EEG) metrics such as non-rapid eye movement (NREM) spindles and slow oscillations (SO), are altered in individuals with schizophrenia (SCZ). However, beyond group-level analyses which treat all patients as a unitary set, the extent to which NREM deficits vary among patients is unclear, as are their relationships to other sources of heterogeneity including clinical factors, illness duration and ageing, cognitive profiles and medication regimens. Using newly collected high density sleep EEG data on 103 individuals with SCZ and 68 controls, we first sought to replicate our previously reported (Kozhemiako et. al, 2022) group-level mean differences between patients and controls (original *N*=130). Then in the combined sample (*N*=301 including 175 patients), we characterized patient-to-patient variability in NREM neurophysiology.

**Results:** We replicated all group-level mean differences and confirmed the high accuracy of our predictive model (Area Under the ROC Curve, AUC = 0.93 for diagnosis). Compared to controls, patients showed significantly increased between-individual variability across many (26%) sleep metrics, with patterns only partially recapitulating those for group-level mean differences. Although multiple clinical and cognitive factors were associated with NREM metrics including spindle density, collectively they did not account for much of the general increase in patient-to-patient variability. Medication regimen was a greater (albeit still partial) contributor to variability, although original group mean differences persisted after controlling for medications. Some sleep metrics including fast spindle density showed exaggerated age-related effects in SCZ, and patients exhibited older predicted biological ages based on an independent model of ageing and the sleep EEG.

**Conclusion:** We demonstrated robust and replicable alterations in sleep neurophysiology in individuals with SCZ and highlighted distinct patterns of effects contrasting between-group means versus within-group variances. We further documented and controlled for a major effect of medication use, and pointed to greater age-related change in NREM sleep in patients. That increased NREM heterogeneity was not explained by standard clinical or cognitive patient assessments suggests the sleep EEG provides novel, nonredundant information to support the goals of personalized medicine. Collectively, our results point to a spectrum of NREM sleep deficits among SCZ patients that can be measured objectively and at scale, and that may offer a unique window on the etiological and genetic diversity that underlies SCZ risk, treatment response and prognosis.

## INTRODUCTION

Schizophrenia (SCZ) is a heterogeneous neuropsychiatric disorder characterized by variable combinations of positive and negative symptoms and cognitive deficits. It is highly heritable and polygenic (International Schizophrenia Consortium et al., 2009; Trubetskoy et al., 2022) and has a substantial impact on affected individuals and their caregivers (Stanley et al., 2017). One of the most urgent tasks in SCZ research is the identification of objective biomarkers for neurobiological deficits to aid in diagnostics, prognostics, patient stratifications, and to guide the novel therapeutic approaches. A growing body of literature (Bagautdinova et al., 2023; Chan et al., 2017; Lai et al., 2022), including our own findings (Kozhemiako et al., 2022), points to sleep neurophysiology as providing a rich array of putative electroencephalography (EEG) biomarkers with robust and replicable group differences between SCZ patients and healthy controls. Notably, we previously showed that sleep-based biomarkers were largely independent of wake EEG metrics from event-related potential paradigms (Erickson et al., 2016; Freedman et al., 2020; Thuné et al., 2016) measured in the same individuals, suggesting the sleep EEG offers unique information about neural underpinnings of SCZ.

However, a biomarker that only tracks with current diagnostic status is arguably of limited value. For a heterogeneous disease such as SCZ, biomarkers that support clinically meaningful stratification of patients, for example by etiology (neurobiological deficits) or likelihood of treatment response, are much needed. We posited that person-to-person variability in sleep micro-architecture may index etiologically relevant heterogeneity among patients. As group-level mean differences tend to reflect only commonalities among patients, biomarkers exhibiting increased patient-specific variability may be more likely to reflect distinct continua of risk or heterogeneous subtypes of disease pathophysiology.

In this study, we therefore searched for sleep-architecture metrics showing increased between-person variability in patients, alongside the standard assessment of differences in group means. We next related the sleep metrics to other measures known to vary among patients, namely clinical symptoms, cognitive deficits and illness duration. Whereas an emerging literature points to replicable group-level alterations in sleep physiology in SCZ (Lai et al., 2022), the broader landscape – of heterogeneity among SCZ patients as well as specificity across other neurological and psychiatric diseases – is less well charted. A recent review focused on sleep spindles, symptomatology and cognitive deficits in SCZ concluded that small sample sizes and inconsistent methodologies led to a high risk of bias and deterred strong conclusions (Au and Harvey, 2020). Despite some support for associations between spindles and attention/cognitive processing speed in patients and the role of spindles in memory consolidation (Manoach and Stickgold, 2019), robust connections between NREM sleep and SCZ symptomatology have not been well established.

Patient-to-patient variation in sleep architecture may also be attributable to different medication regimens. Psychoactive drugs significantly impact sleep patterns as well as the sleep EEG (Leong et al., 2022), although associations can be complex: they can normalize some aspects of sleep in SCZ but disrupt others (Krystal et al., 2008). While studies of unmedicated SCZ patients have established that sleep abnormalities (notably, spindle deficits) persist independent of medication (Manoach et al., 2014), recent reviews have pointed to the need for research to better characterize the roles of medications in generating sleep alterations (Ferrarelli, 2023; Kaskie and Ferrarelli, 2020). In particular, larger sample sizes – such as offered by the current study – will be necessary to resolve the impact of different medications on sleep architecture in patients and to disentangle it from underlying disease-associated signatures.

Finally, beyond illness duration per se, variation among patients may reflect the differential effects of ageing. A substantial imaging literature has pointed to accelerated brain ageing in SCZ (Constantinides et al., 2023; Kaufmann et al., 2019), which may in part account for its increased burden of age-related morbidity and premature mortality (Hjorthøj et al., 2017). The NREM EEG changes profoundly across typical development (Purcell et al., 2017) and delayed or accelerated patterns have also been shown to predict diverse pathologies (Kozhemiako et al., 2023; Sun et al., 2019). We therefore also analyzed the sleep EEG data through the lens of biological/brain age prediction to address the hypothesis of accelerated brain ageing in SCZ.

Here we report on our ongoing Global Research Initiative of Neurophysiology on Schizophrenia (GRINS) study, in particular the independent second wave of *N*=103 SCZ patients and *N*=68 controls with overnight high-density EEG and extensive clinical, cognitive and demographic data. To establish whether sleep macro-and micro-architecture provide a robust and novel window on SCZ heterogeneity, we first sought to replicate our previously reported group-level mean differences in sleep physiology (Kozhemiako et al., 2022). Subsequently, in the combined sample (*N*=301) we determined whether between-individual variability in the same set of metrics was altered in SCZ. Finally, we investigated whether differences could be explained by measured clinical and cognitive factors, illness duration and ageing, or medication use.

## RESULTS

GRINS wave 2 comprised *N*=103 SCZ and *N*=68 control (CTR) individuals, newly collected under the same protocol as wave 1 (see (Kozhemiako et al., 2022) for details). Demographic and sleep variables are given in **Table 1**, for each wave independently as well as the combined sample (total *N*=175 SCZ, *N*=126 CTR). Wave 2 closely resembled wave 1 for primary demographic, clinical, cognitive and sleep variables. Wave 2 controls were slightly older with less N3 sleep compared to wave 1 controls (*p* < 0.01) while wave 1 controls were slightly younger (p<0.01), and in the combined sample there were no significant case-control differences in either age or N3 duration.

**Table 1.** Key demographic and clinical variables stratified by wave. * – differences between SCZ and CTR groups (p < 0.01) ^#^ – differences between Wave 1 and Wave 2 within the diagnostic group (p < 0.01) ª – such scores are considered to represent mild symptoms

| *Characteristics* | *Wave 1* |  | *Wave 2* |  | *Combined* |  |
| --- | --- | --- | --- | --- | --- | --- |
|  | *SCZ* | *CTR* | *SCZ* | *CTR* | *SCZ* | *CTR* |
| *N* | 72 | 58 | 103 | 68 | 175 | 126 |
| *Sex, females (%)* | 25 (35%) | 22 (31%) | 49 (48%) | 31 (30%) | 74 (42%) | 53 (30%) |
| *Age, years* | 35±7 | **32±6.3** | 34±7.5 | **36±6<sup>#</sup>** | 35±7.3 | 34±6.5 |
| *Maternal education, higher than middle school (%)* | 30 (42%) | 14 (24%) | 29 (28%) | 21 (31%) | 59 (34%) | 35 (28%) |
| *Paternal education, higher than middle school (%)* | 31 (43%) | 22 (38%) | 37 (36%) | 21 (31%) | 68 (39%) | 43 (34%) |
| *PANSS positive* | 15±5.8 <sup>a</sup> |  | 17±6.3 <sup>a</sup> |  | 16±6.1 <sup>a</sup> |  |
| *PANSS negative* | 16±5.7 <sup>a</sup> |  | 18±6.7 <sup>a</sup> |  | 17±6.4 <sup>a</sup> |  |
| *PANSS general* | 33±9 <sup>a</sup> |  | 35±10.9 <sup>a</sup> |  | 34±10.1 <sup>a</sup> |  |
| *SCZ duration, years* | 12±7 |  | 10±6.6 |  | 11±7 |  |
| *MCCB total* | 43±9.9 |  | 44±8.7 |  | 44±9.2 |  |
| *Equivalent antipsychotic dose, mg* | 612±262.8 |  | 676±244.6 |  | 649±253.6 |  |
| *Time in bed, mins* | **532±55.7\*** | **467±44.5** | **540±43.9\*** | **456±27.7** | **537±47.8\*** | **461±36.8** |
| *Total sleep time, mins* | 378±95.1 | 382±63.1 | 398±84.2 | 377±48.1 | 390±88.9 | 376±59.4 |
| *Wake after sleep onset, mins* | 65±44.1 | 49±34.4 | 70±46.8 | 55±39.7 | **67±43.9\*** | **52±37.4** |
| *Sleep efficiency (TST/TIB)* | **73±14.1\*** | **82±10.9** | **74±13.8\*** | **83±9.5** | **74±13.8\*** | **82±10.1** |
| *Sleep maintenance efficiency (TST/start-end of sleep)* | 81±13.8 | 86±10.2 | 85±9.8 | 87±8.7 | **83±11.9\*** | **87±9.4** |
| *N1 duration, mins (%)* | 42±26.2 (11%) | 32±17.3 (9%) | 33±16.8 (9%) | 33±16.5 (9%) | 37±20.4 (10%) | 32±16.8 (9%) |
| *N2 duration, mins (%)* | 186±69.8 (50%) | 204±48.2 (53%) | 209±72.2 (53%) | 217±37.7 (57%) | 201±73.5 (52%) | 211±43.2 (56%) |
| *N3 duration, mins (%)* | 76±49.8 (21%) | **80±31** (21%) | 72±48.4 (18%) | **60±26.5<sup>#</sup>** (16%) | 74±48.9 (19%) | 69±30.3 (18%) |
| *REM duration, mins (%)* | 66±37.6 (17%) | 65±25.1 (17%) | 76±34.4 (19%) | 66±22.9 (18%) | 72±36.1 (18%) | 65±23.8 (17%) |
| *REM latency, mins* | 121±55.2 | 115±59 | 112±57.4 | 97±38.6 | 116±56.5 | 105±49.8 |
| *Number of cycles* | 4±1.4 | 4±0.7 | 4±1.2 | 4±0.9 | 4±1.3 | 4±0.8 |
| *Cycle duration, mins* | 99±28.3 | 104±21.9 | 97±25.2 | 93±18.5 | 98±26.4 | 98±20.8 |

Consistent with wave 1 findings, in the combined sample the SCZ group showed increased time in bed (∼1.5 hours, *p* = 9 × 10^−12^) and decreased sleep efficiency (effect size [e.s.]= −0.95 standard deviation units (SD), *p* = 7 × 10^−6^), primarily driven by longer sleep onset latency. The SCZ group comprised an inpatient sample with a hospital-imposed routine: whereas this implicitly controlled certain factors such as light exposure among patients, it also precluded naturalistic assessment of sleep/wake rhythms and circadian factors. Cases and controls were otherwise well-matched in terms of total sleep time and all stage-specific sleep duration measures (**Table 1**).

### Replication of sleep alterations reported in wave 1

Following our previous work (Kozhemiako et al., 2022), we computed a battery of metrics mainly focused on the NREM sleep EEG. Specifically, we tested 26 unique classes of metric across seven domains (**Table 2**): macro-architecture, fast (FS) and slow (SS) spindles, slow oscillations (SO), spindle/SO coupling, spectral power (PSD) and functional connectivity (PSI, phase slope index summarized channel-wise). Some metrics were calculated for each of the 57 EEG channels and across a range of frequencies, in total yielding 4,746 variables. To address multiple testing, in addition to wave 2 providing an independent replication sample, we further used cluster-based permutation to control false positive rates across channels and frequencies, in the initial replication analysis as well as subsequent combined sample analyses. For the latter analyses, we also applied false discovery rate correction within each of the seven domains.

**Table 2.** NREM metrics tested in Wave 1 (Kozhemiako et al., 2022) and tested for replication in Wave 2.

| Domain | # of metrics | Stratifications | # of tests | Multiple comparison correction for replication | # PCs extracted |
| --- | --- | --- | --- | --- | --- |
| *Macro-architecture* | 9 | $\times 4$ stages (for 2 metrics) | 15 | None ($p$-value threshold $< 0.01$) | - |
| *Slow oscillations (SO)* | 5 | $\times 57$ channels | 285 | Cluster-based across channels | 3 |
| *Slow Spindles (SS)* | 6 | $\times 57$ channels | 342 | Cluster-based across channels | 4 |
| *Fast Spindles (FS)* | 6 | $\times 57$ channels | 342 | Cluster-based across channels | |
| *Spindle/SO coupling* | 4 | $\times 2$ spindle frequencies<br>$\times 57$ channels | 456 | Cluster-based across channels | - |
| *Channel-wise connectivity (PSI)* | 1 | $\times 18$ frequencies<br>$\times 57$ channels | 1026 | Cluster-based across channels & frequencies | 3 |
| *Spectral power (PSD)* | 1 | $\times 40$ frequencies<br>$\times 57$ channels | 2280 | Cluster-based across channels & frequencies | 2 |

All previously reported wave 1 sleep EEG group differences replicated in wave 2, based on a significant metric-level result following correction for multiple comparisons and a similar direction of effect (**Figure 1A**). For example, in wave 1 fast spindle (FS) density at C2 was reduced by 28% in SCZ (2.2 vs 3.1 spindles per minute in CTR) and by 30 % (2.3 vs 3.3) in wave 2. Spatial patterns of group differences were broadly congruent between waves, for example, the reduction in slow spindle (SS) density in posterior channels (**Figure 1A, Figure S1**).

**Figure 1.**
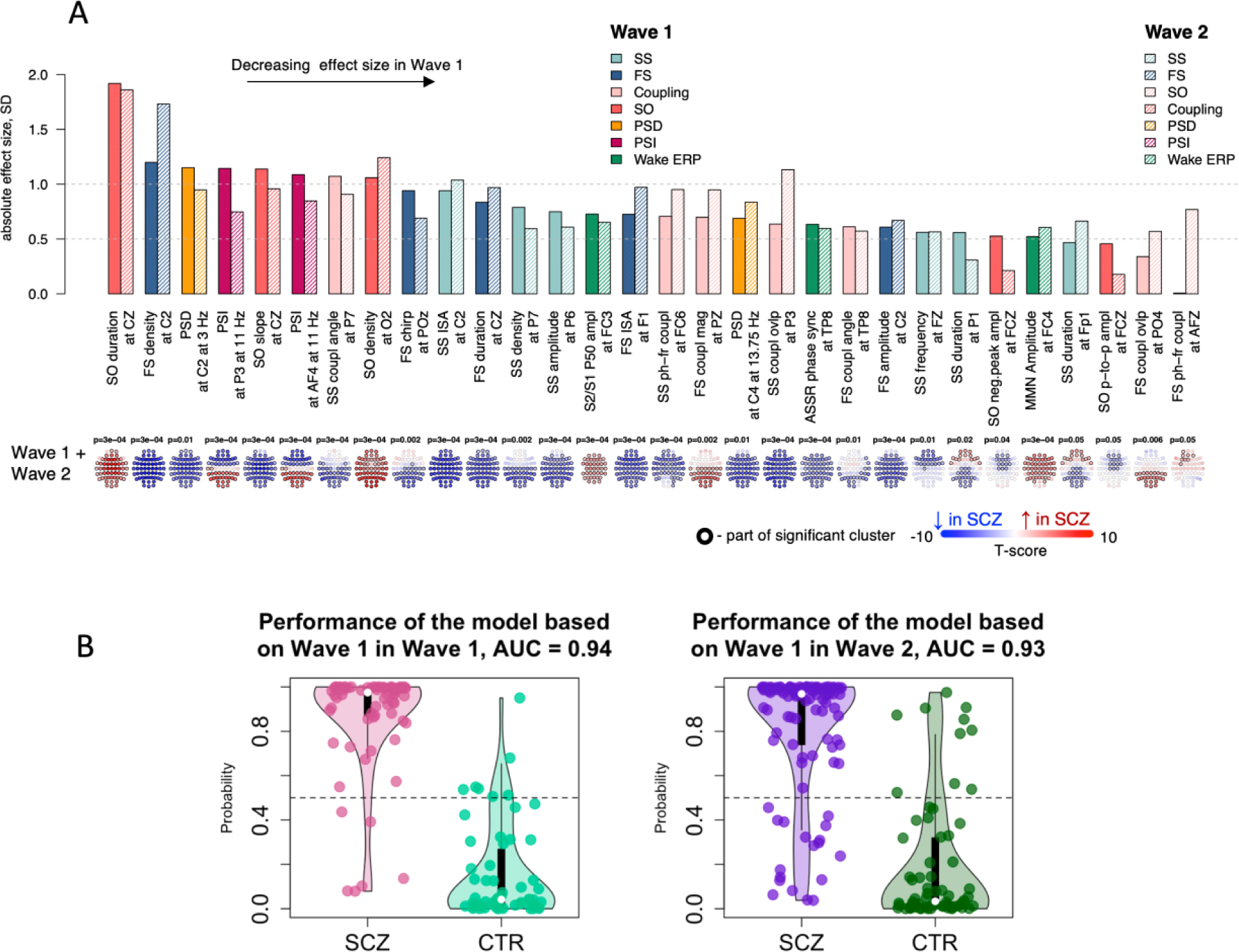
Replication of wave 1 sleep neurophysiology alterations in wave 2 of the GRINS cohort. **A –** absolute effect sizes of group-level mean differences in wave 1 and wave 2 respectively, ordered from highest to the lowest (note: the directions of effect were consistent between wave 1 and wave 2). Metrics with significant combined sample SCZ-CTR clusters are displayed as topoplots; the EEG channel with the highest absolute t-score was selected for top bar plot. Metrics are ordered from left to right based on decreasing effect size in wave 1. **B –** SCZ-CTR classification in wave 2 (Right) using a predictive model derived from wave 1 data (Left).

As expected from the increased sample size, we detected novel associations in the combined sample which did not pass our stringent significance criterion in wave 1 (cluster statistics are summarized in **Table S1**). These included the SCZ group showing i) longer frontal and shorter posterior SS duration (two clusters with maximum effects at Fp1 e.s. = 0.55 SD, *p* = 7×10^−4^ and P1 e.s. = −0.41 SD, *p* = 0.003 respectively), ii) decreased SS frequency in frontal channels (at FZ e.s. = −0.57 SD, *p* = 2×10^−5^), iii) increased FS/SO coupling magnitude and overlap in posterior channels (at PZ e.s. = 0.86 SD, *p* = 5 × 10^−8^ and at PO4 e.s. = 0.51 SD, *p* = 8 × 10^−5^ respectively). Only a single metric showed a qualitative difference in the evidence for statistical association between waves, namely SO-phase FS-frequency modulation, which was increased in frontal channels only in wave 2 (e.g. at AFZ, e.s. = 0.34 SD, *p*=0.001).

In addition to replication of individual metrics, we evaluated the performance of our NREM-based prediction model to classify diagnostic status. This logistic regression model was trained on wave 1 data only, using 12 principal components (PC) summarizing spindle, SO, spectral power and functional connectivity metrics (see **Table 2** and Kozhemiako et al., 2022 for details). Projecting wave 2 individuals into the predefined 12-dimensional PC space, classification performance in wave 2 was effectively identical to wave 1, with an Area Under the ROC Curve (AUC) value of 0.93 versus 0.94 from the original wave 1 analysis (**Figure 1B**).

### Spindle density deficits in SCZ depend on temporal coupling with slow oscillations (SO)

Having replicated group-level mean differences, subsequent analyses were performed using the combined (*N*=301) sample. In replication and combined analyses (**Figure 1**), SCZ patients showed reductions in fast and slow spindles, as well as differences in the rate of and temporal coupling with SO. To further characterize altered spindle/SO coupling in SCZ, we computed the density (count per minute) of SO-coupled and SO-uncoupled spindles separately. Approximately 25% to 45% of spindles (depending on channel and spindle class) overlapped (“coupled” with) a detected SO. Case/control reductions in overall spindle density reflected qualitatively different effects for coupled and uncoupled spindles (**Figure 2**). For both fast and slow spindles, the overall reductions in spindle densities were largely driven by fewer SO-uncoupled spindles in patients. In contrast, controlling for SO-uncoupled spindle density, SO-coupled spindles either showed no group differences (slow spindles) or even a significant increase in patients (fast occipital spindles). This latter result is consistent with the previous significant increase seen for the fast spindle-SO overlap statistic: in patients there are fewer fast spindles overall, although some types (specifically, fast occipital SO-coupled spindles) are relatively over-represented, underscoring that topographical and temporal contexts are important to consider when evaluating spindle activity.

**Figure 2.**
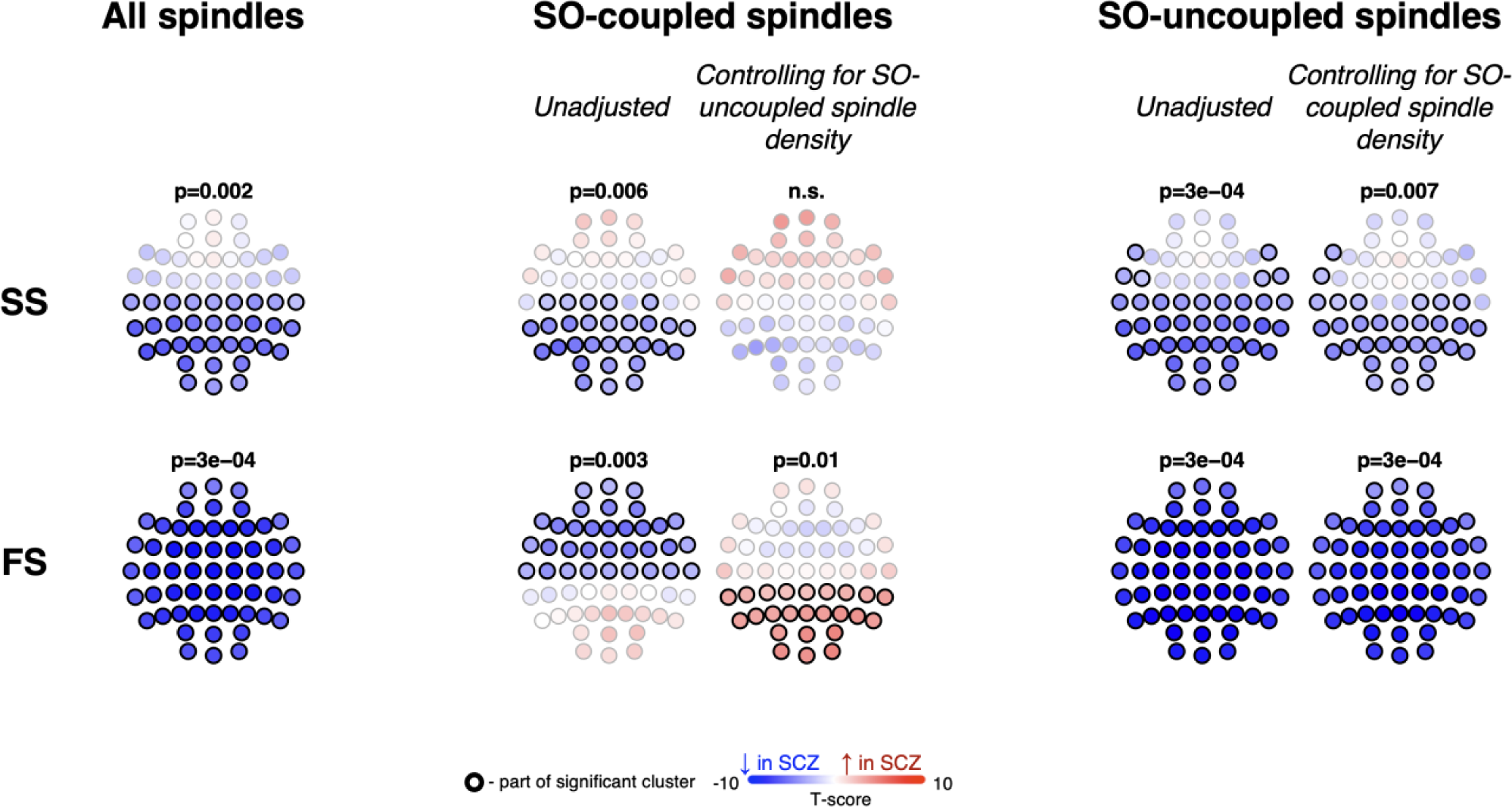
Spindle density deficits in SCZ depend on SO-coupling status. The topoplots represent the group differences between SCZ and CTR in spindle density computed based on all, coupled with SOs and uncoupled with SOs spindles. The first row displays the results for slow spindles (SS), while the second row shows the results for fast spindles (FS).

### Greater between-individual variability among SCZ patients across diverse sleep metrics

We next focused on between-individual variability for the sleep metrics in **Table 2**. Of these, 1,232 (26%) exhibited nominally (*p* < 0.05) significant differences, based on Bartlett’s test comparing the between-individual variances within the SCZ group versus within the CTR group, after removing outliers and adjusting for the effects of age and sex. Of note, all but 15 significant tests pointed to higher variability in the SCZ group (**Figure 3A**).

**Figure 3.**
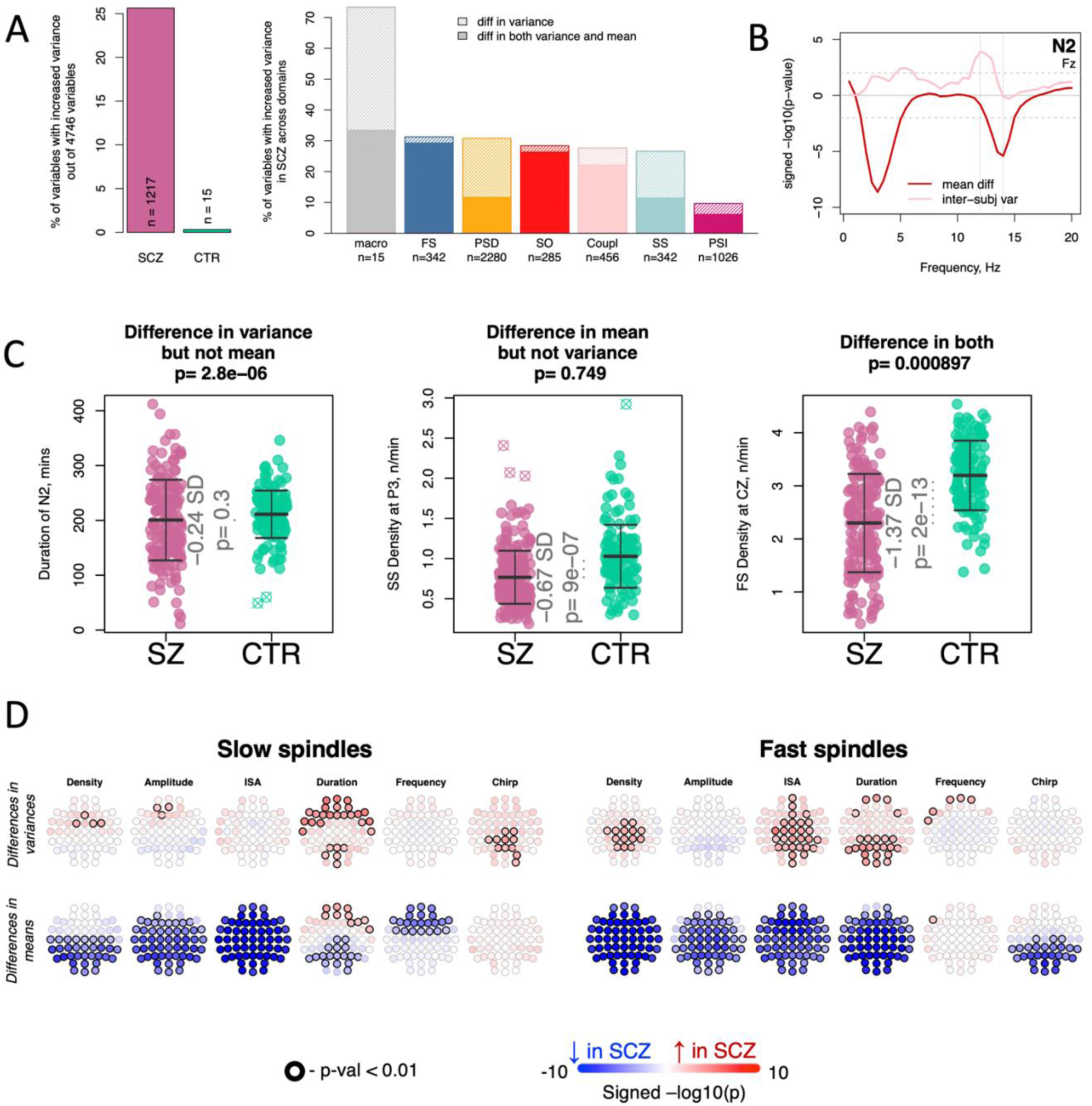
Increased variability in the SCZ group across multiple sleep estimates. **A –** Left : the percentage of sleep variables with significantly (p < 0.05) increased variability in SCZ vs CTR; Right: the percentage of sleep variables with increased variability (light shade) and those with both increased variability and altered means (dark shade) across seven domains. **B –** visualization of differences in means and inter-individual variances of spectral power at FZ across frequencies during the NREM2 stage; vertical lines represent 12 and 14 Hz. **C –** examples of sleep variables with the difference in variance (Bartlett’s test p-value in the title with effects of age and sex regressed out prior statistical comparison), but not in means (effect size and p-value from logistic regression controlling for age and sex inside the graph); in mean, but not in variance; and in both mean and variance. **D** – topoplots illustrate the distinct profiles of between-group differences in variance (top row) versus mean (bottom row) all channels for FS and SS metrics

Macro-architecture metrics expressed some of the largest variance increases in SCZ. For example, whereas N2 duration showed no significant difference in means (201 vs 209 minutes, *p* = 0.26), cases had a markedly higher standard deviation (SD) of 71, versus 47 for controls (Bartlett *p* = 3 × 10^−6^). Total sleep time, N3 and REM duration likewise showed similar increases in variability in SCZ. This signature of increased SCZ variability was further observed across all domains of NREM micro-architecture. Whereas for some domains, metrics exhibiting increased variance among SCZ individuals almost always showed concurrent significant mean differences (primarily FS, SO and spindle/SO coupling), metrics in other domains also exhibited variance differences in the absence of corresponding group difference in means (primarily macro-architecture, SS and PSD domains, **Fig 3A**).

**Figure 3B** shows SCZ-CTR differences for stage N2 spectral power (PSD) at a representative frontal channel (Fz): whereas slower sigma frequencies (∼11.5 Hz) showed different variances (higher in SCZ) but equivalent means, faster sigma frequencies (13-16 Hz) showed the opposite pattern, of similar variances but a significantly lower mean in SCZ. As a second illustration of these divergent effects, **Figure 3C** shows three exemplar metrics with qualitatively distinct alterations in SCZ: in variance only (N2 duration), in means only (posterior SS density), or in both (central FS density). Similarly, primary spindle metrics showed distinct patterns for group differences in variances versus means (**Figure 3D**).

In principle, increased inter-individual variability in SCZ is consistent with the hypothesis that the sleep EEG stratifies patients into discrete subtypes. However, we did not find evidence of distinct patient-specific clusters readily emerging from exploratory analyses using either principal component or UMAP (uniform manifold approximation and projection) dimension reductions, whether based on metrics showing increased variances or with altered means in the SCZ group (**Figure S2**).

### Patient heterogeneity in clinical and cognitive factors

We next asked whether the increased NREM heterogeneity among SCZ patients could be accounted for by patient-to-patient differences in illness duration, symptoms or cognitive deficits. We created new sets of sleep metrics for patients, that adjusted for either clinical (illness duration, total antipsychotics dosage and PANSS scores) or cognitive variables (MST overnight improvement and morning test performance, MCCB scores), using residuals from patient-only linear regressions that fit each sleep metric jointly on all clinical or cognitive variables, thereby removing the between-patient variability explained by these factors. Repeating the Bartlett tests, the SCZ group nonetheless continued to exhibit significantly higher variabilities for the majority of the previously reported 1217 metrics: 965 or 870, controlling for clinical or cognitive variables respectively (**Figures 4A & 4B**).

**Figure 4.**
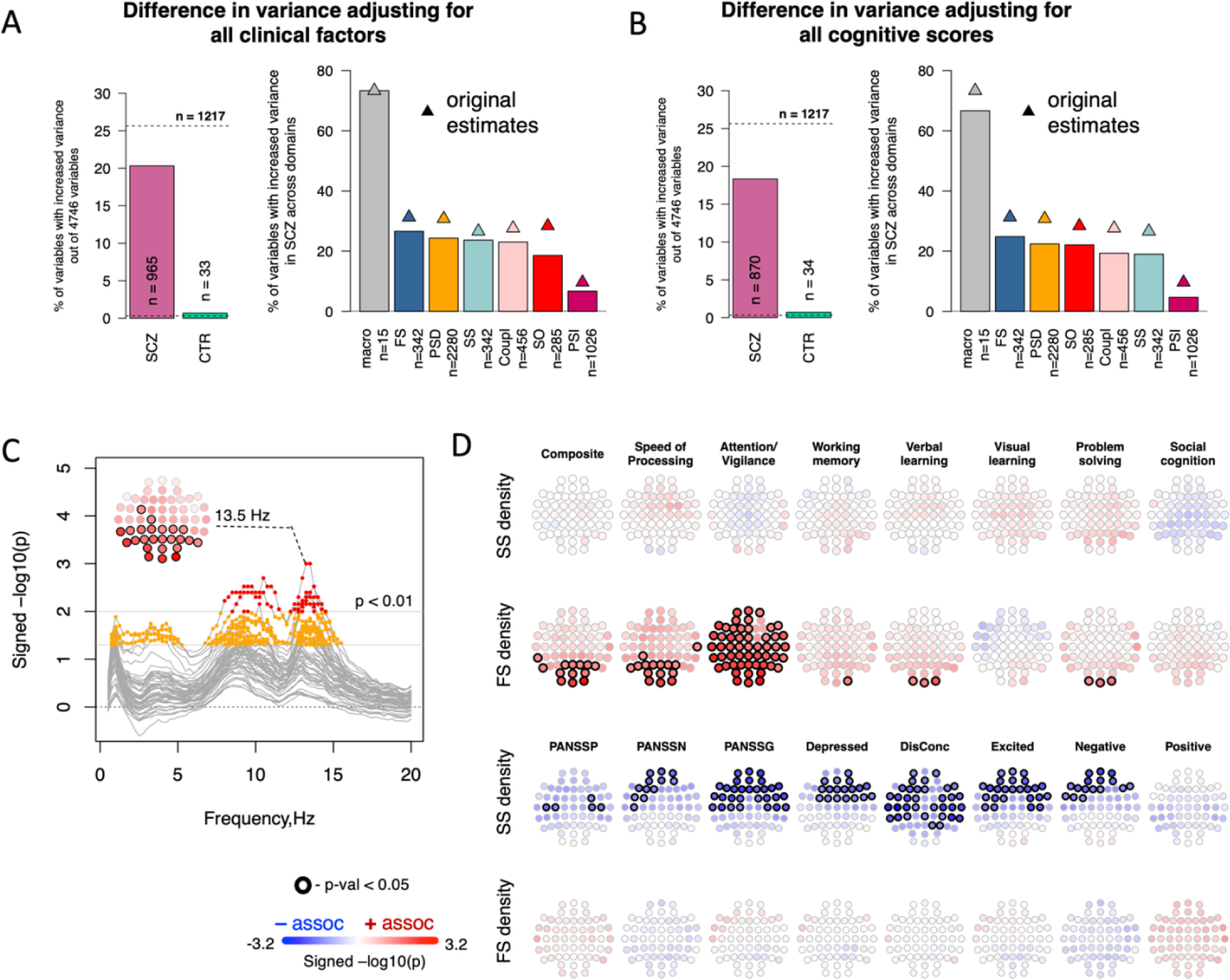
Sleep EEG associations with clinical and cognitive factors. **A –** the left bar plot shows the percentage of all sleep variables still exhibiting higher variance in either the SCZ or CTR group after the effects of all clinical factors were simultaneously regressed out (for the SCZ group only), compared to the original estimates (horizontal dashed line); the right bar plot is similar but is stratified by domain (the denominator for percentages is the total number of variables in the domain); original estimates are marked by triangles. **B –** same as A but controlling for the cognitive (instead of clinical) variables. **C** – the graph illustrates the association between the total MCCB score and spectral power across all channels (lines) in the frequency range of 0.5-20 Hz. Channels and frequencies displaying association above the nominal significance threshold of p<0.05 are highlighted in orange and with p <0.01 in red. **D** – topoplots represent channels where the association between spindle density and total MCCB score (top two rows) or PANSS scores (bottom two rows) were nominally significant (p <0.05).

The persistence of increased NREM variability in SCZ suggests that objective sleep biomarkers may capture disease-relevant factors that are not otherwise well-characterized by existing clinical and cognitive measures. Nonetheless, it does not imply that NREM sleep metrics were unrelated to clinical and cognitive factors in patients. Although a comprehensive examination is beyond the scope of this manuscript, we performed a series of secondary analyses focused on patient MCCB and PANSS scores in relation to N2 spectral power and spindle density. Higher MCCB composite scores were associated with increased occipital and parietal N2 sigma-range power (maximal at 13.5 Hz, *p* < 0.01, **Figure 4C**). Consistent with this, MCCB composite scores were also associated with increased fast spindle density (**Figure 4D**). Considering the separate MCCB domains, this effect was most pronounced for Attention/Vigilance and Speed of Processing domains and was restricted to fast spindles. In contrast, clinical measures (three primary PANSS scores and derived five-factor scores (Wallwork et al., 2012)) showed nominal (*p*<0.05) associations with slow spindle densities at multiple central and frontal channels, across most clinical measures, whereby reduced slow spindle activity predicted more severe symptoms (**Figure 4D**).

### NREM sleep predictors of sleep-dependent motor procedural memory in SCZ

Sleep-dependent motor procedural memory (finger tapping motor-sequence task, MST) has previously been associated with greater %N2 in healthy individuals and fast sleep spindle density in SCZ patients (Mylonas et al., 2020a; Walker et al., 2002; Walker and Stickgold, 2004; Wamsley et al., 2012). We observed a robust attenuation of overnight improvement of MST performance in the SCZ group versus controls (percentage, e.s. = −0.99, p=4×10^−9^, **Figure S3**), while learning rate during training was not different between the groups (*p* > 0.05), consistent with prior reports (Manoach et al., 2010, 2004; Wamsley et al., 2012). However, neither %N2 nor sleep spindle density were associated with overnight MST performance in the SCZ group.

### Medication effects

Medication use showed an array of robust effects on both macro-and micro-architecture sleep metrics (**Figures 5 & S4)**. Each patient’s medication use at the time of the sleep study was encoded as a binary vector of nine medications (six antipsychotics and three categories of adjunct drugs, each used by at least 10 patients) and entered into a case-only linear model to predict each sleep metric, controlling for age and sex. Among antipsychotic medications, olanzapine (*N*=81 patients) and clozapine (*N*=22) affected sleep metrics the most (**Figures 5A, 5B**), albeit sometimes in qualitatively different ways. For example, whereas olanzapine use was associated with decreased N2 proportion (by −10 % compared to patients not taking olanzapine) and increased N3 (+7%), clozapine use had the opposite pattern of increased N2 (+14%) and decreased N3 (−9%); likewise, they had opposite effects on SS coupling overlap (olanzapine: max *t*-score=-3.4, cluster *p* = 0.003; clozapine: max *t*-score=4.2, cluster *p* = 0.001).

**Figure 5.**
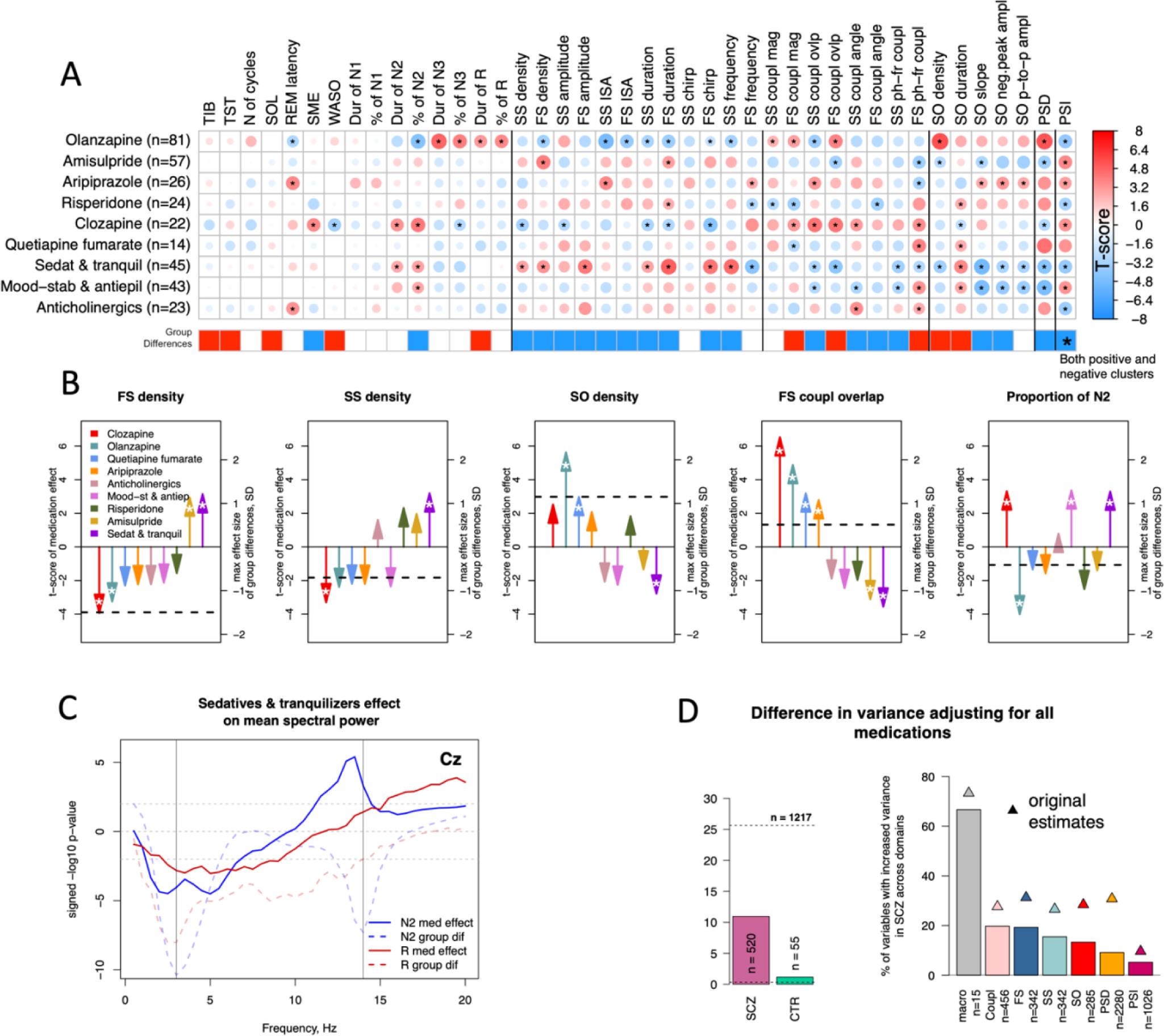
Sleep EEG associations with medication use in SCZ. **A –** the matrix shows t-scores from linear regressions (controlling for age and sex) between sleep features and binarized medication use in SCZ (where each medication is included separately). For multi-channel variables, the largest t-score among the channels is presented and stars mark associations with p < 0.01. The horizontal bar below the matrix illustrates SCZ-CTR differences in the corresponding variable: blue = decrease, red =increase, white =no difference; **B –** examples of medication effects on FS density, SS density, SO density, FS coupling magnitude and N2 proportion. Each arrow indicates an effect (t-score) of a certain medication on a sleep variable from a multiple linear regression where all medication groups were included as well as age and sex. Nominal significance (p<0.05) is marked by a white star inside the arrows. For multi-channel metrics, the largest effect across all channels is presented. The horizontal dashed line indicates the effect size of the group difference between SCZ and CTR in the corresponding metric (at the channel with the largest effect size for multi-channel variables). **C –** sedatives and tranquilizers effect on spectral power in N2 and REM across frequencies (solid lines) in comparison to SCZ-CTR differences (dashed lines) at Cz channel **D –** left bar plot shows the percentage of all sleep variables still with higher variance in SCZ or CTR group after effects of all common medications have been simultaneously regressed out for SCZ group, compared to the original estimates (horizontal dashed line); the right bar plot is similar but stratified by domain (denominator for percentages is the number of variables in the domain); original estimates are marked by triangles.

Among adjunct medications, sedative and tranquilizer use (*N*=45) had the most marked effects on sleep micro-architecture, impacting multiple spindle (e.g. increased FS duration, *p* = 4×10^−4^, **Figure S4**) and SO characteristics (e.g. altered slopes, *p* = 3×10^−4^). Sedative and tranquilizer use was also associated with decreased 1 - 7 Hz power during N2 (cluster *p* = 0.009, **Figure 5 C**). The effect on 1-7 Hz power was also present during REM sleep, as well as with mood stabilizer and antiepileptic use, albeit with an attenuated effect.

Given the variability in patterns of medication use, combined with the marked and sometimes divergent associations with the sleep EEG, we posited that medication use would account for some of the increased variability in sleep metrics found in SCZ. Adopting the same approach as above (adjusting for medication use in patients prior to testing for group differences in variance using Bartlett’s test), we observed a greater – but still only partial – drop in the number of metrics showing increased variance in SCZ. The largest declines were in the domain of spectral power, from approximately 30% to 10% of all metrics that are significantly more variable, with an expected rate of 5% given the nominal significance threshold of *p* = 0.05, **Figure 5D**). Nonetheless, across all domains, 520 metrics – a rate far greater than expected by chance – still showed greater variance among SCZ patients, compared to only 55 metrics with a reduced variance in controls.

### Accelerated age-related NREM alterations in SCZ

Finally, we asked whether SCZ patients have greater sensitivity to other factors known to influence sleep in the general population – in particular, age and sex (note: these were statistically controlled in the above analyses). In exploratory models including interaction terms allowing the effects of age and sex to vary between cases and controls, the most prominent interaction involved age and FS density across multiple channels, such that the SCZ group showed greater age-related declines (maximal effect at F5, interaction *p* = 4 × 10^−4^; 46 channels had an interaction *p* < 0.01, **Figure 6A**). This effect was specific to FS: SS density interaction terms were nonsignificant (*p* > 0.05) for all channels. Sex did not appear to modify case/control differences in either FS or SS (all *p* > 0.05). Although spindle density decreases in older adults (Purcell et al., 2017), in our middle-aged (median age 34, IQR 30.5 – 39) sample FS density was largely independent of age in controls (e.g. *r* = −0.03, *p* = 0.76 at F5). In contrast, the SCZ group showed a pronounced age-related reduction in FS density (*r* = −0.35, *p* = 4 × 10^−6^), consistent with an accelerated ageing effect among individuals with SCZ (**Figure 6A**).

**Figure 6.**
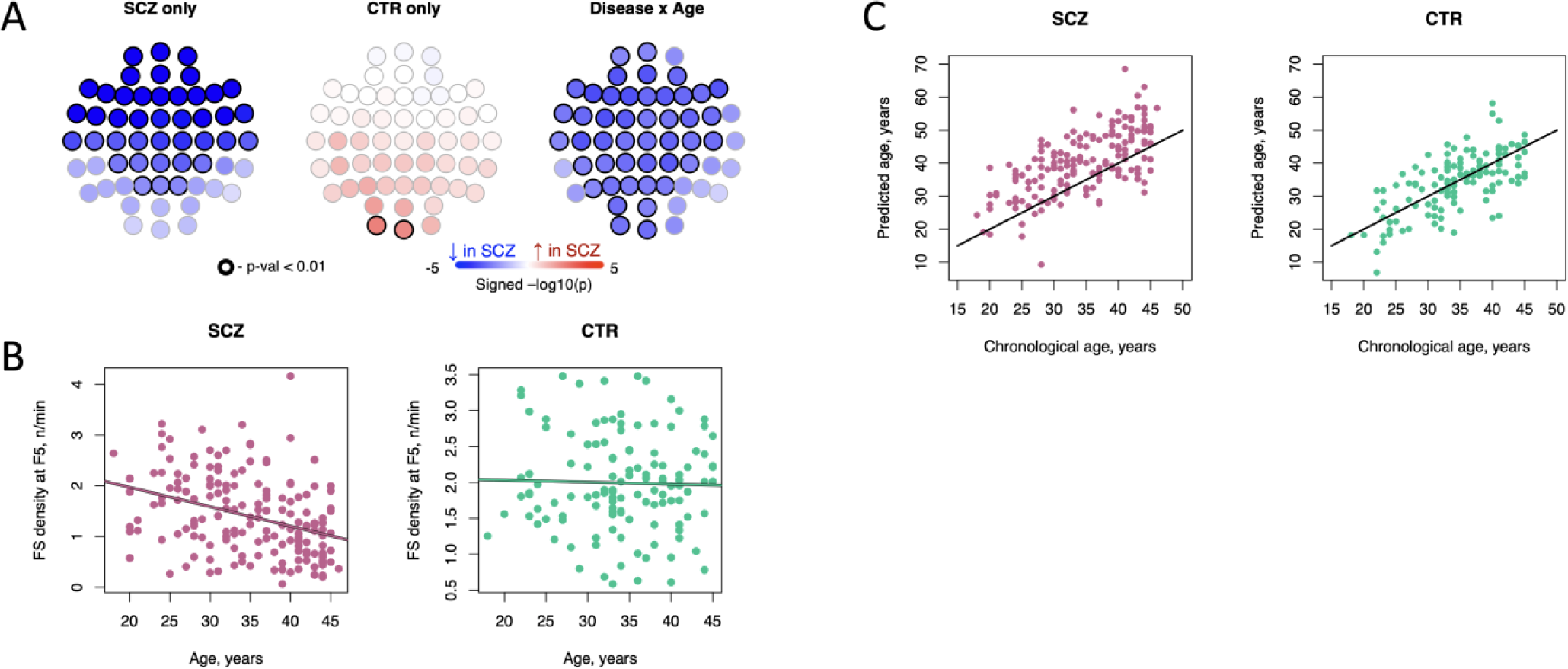
Fast spindle density shows differential age-related decline in SCZ. **A –** the topoplots show the associations between FS density and age, in either a case-only (left plot) or control-only (middle plot) analysis, and the interaction P-values from the joint models (right plot); associations P < 0.01 are marked by dark circles; **B** – scatter plots showing FS density at F5 as a function of age separately in cases and controls. **C** – scatter plots showing predicted and observed ages separately in cases and controls; age prediction was based on a modified version of the model described by Sun et al (2019).

Key macro-architecture metrics including TST also showed significant age × disease interactions. Although patients were well-matched to controls in terms of age (*t*-test and Bartlett test *p* = 0.68 and *p* = 0.19, comparing means and variances in age between groups, respectively), patients showed a highly significant age-related reduction in TST (*r* = −0.32, *p* = 1 × 10^−5^), whereas TST and age were unrelated in controls (*p* = 0.57), yielding a significant age × disease interaction (*p* = 0.003). Controlling for illness duration and age-at-onset did not alter the TST-age association in patients and neither term was associated with TST conditional on age. Likewise, patient FS density was not associated with either illness duration or age-at-onset when controlling for chronological age.

To more directly address the question of accelerated ageing, we used an independently developed model to predict so-called brain age from the sleep EEG ((Sun et al., 2019), see Methods for details). Whereas predicted and chronological ages were similarly correlated in patients (*r* = 0.65) and controls (*r* = 0.67), predictions for patients were on average 5.8 years greater than their chronological age (*p* < 10^−15^ one-sample *t-*test),whereas in controls predicted and chronological ages had similar means (−0.1 year difference, *p* = 0.85). Consequently, the predicted age difference (PAD = predicted age – chronological age) was significantly higher in patients compared to controls (*p* = 10^−12^, also covarying for chronological age and sex), consistent with accelerated ageing. However, the PAD did not show greater variability among SCZ patients compared to controls (Bartlett *p* = 0.27). In patients, PAD was not associated with duration of illness (*p* = 0.19 in a model controlling for chronological age and sex).

In controls, the mean PAD varied between males (+1.17 years) and females (−1.87 years) significantly (*p* = 0.007). In contrast, the PAD in male (+6.4 years) and female (+5.0 years) patients were not significantly different from each other (*p* = 0.24). Furthermore, the PAD was not associated with clinical (PANSS) or cognitive (MCCB) scores (*p* > 0.05), although it was associated with olanzapine use, such that patients using olanzapine (*N*=81) had significantly (*p* = 2×10^−8^) older PAD scores (+8.98 years) compared to patients not using olanzapine (*N*=94; mean +3.08 years); this effect remained significant (*p*=1×10^−5^) when jointly controlling for other medications used. Olanzapine use did not account for the original case-control difference in PAD score however, which was present albeit attenuated in both groups (*p* < 10^−15^ for olanzapine users versus controls, compared to *p* = 5×10^−4^ comparing patients not using olanzapine to controls).

## DISCUSSION

In a large high-density EEG study of sleep and SCZ, we found unambiguous support for our previously reported alterations of NREM neurophysiology (Kozhemiako et. al., 2022), observing comparable effect sizes and directions. Our predictive model of diagnostic status, previously trained on only the first wave of data, rendered effectively identical classification accuracy in the independent second wave. Overall, these replicated results implicate not only reduced spindle density as a core feature of SCZ, but also multiple aspects of N2 sleep including spindle morphology, SO features and spindle-SO coupling, as well as differences in spectral power and patterns of functional connectivity.

Independent of group differences in means, patients with SCZ showed significantly increased person-to-person variability for many sleep metrics considered. This echoes previous reports of higher between-individual variability in brain morphology (Alnæs et al., 2019; Brugger and Howes, 2017) and functional connectivity (Gopal et al., 2016) in SCZ, as well as the long-standing nosological discussion of heterogeneity (Tsuang and Faraone, 1995; van Os et al., 2010). Such heterogeneity could be linked to existence of quasi-discrete SCZ subtypes – an hypothesis tested by prior studies attempting to derive subtypes based on symptom profiles or cognitive functioning, but which failed to show clear links to specific neurophysiological mechanisms (Seaton et al., 2001). Although not a primary focus of this study, preliminary analyses did not suggest clearly distinct patient sub-groups on the basis of NREM alterations. Rather, our results point to a continua or spectrum of NREM alterations among patients. This spectrum of NREM changes may reflect impairments in multiple neurocircuits, consistent with the polygenic nature of SCZ disease risk that suggests many biological pathways underlying its pathophysiology.

Whereas cognitive deficits are often reported in SCZ (Au and Harvey, 2020; Carruthers et al., 2021), multiple clinical and cognitive factors exhibit significant diversity among patients (Ahmed et al., 2018; Demjaha et al., 2017). Our results suggest a limited connection between such factors and sleep neurophysiology in patients. Increased FS density predicted fewer cognitive deficits (in particular, MCCB attention/vigilance scores) and increased SS density predicted less severe symptoms. However, these associations were generally modest, and the observed variability in clinical and cognitive factors could not account for the increased patient variability in sleep. This is in line with work suggesting that sleep spindles characteristics are poor predictors of general neurocognitive functioning in patients (Baandrup et al., 2019), as well as a recent review that reported little consistency and a lack of robust associations between specific spindle metrics and symptom measures (Au and Harvey, 2020). In earlier work, the use of small samples and diverse analytic and design factors precluded strong conclusions however, either positive or negative.

With regard to MST performance, although we replicated prior findings of impaired sleep-dependent memory consolidation in SCZ (Demanuele et al., 2017; Demirlek and Bora, 2023), previous reports linking overnight MST improvement and spindle density (Mylonas et al., 2020a; Wamsley et al., 2012) were not supported by the current study. Future work will be needed to resolve these apparent discrepancies, whether they reflect purely statistical (type I or type II) errors, or systematic differences in factors such as sample composition, demographics, medication regimens or inpatient versus outpatient settings.

Patterns of medication use induced significant heterogeneity in patient sleep EEG metrics. Although several relatively small studies reported direct, typically acute effects of antipsychotic use on sleep (Arai et al., 2023; Göder et al., 2008; Kluge et al., 2014; Tsekou et al., 2015), our study focused on a naturalistic setting of chronic use in a wide range of antipsychotics, with patients often on multiple medications concurrently. Even though all antipsychotics considered here belonged to the second-generation (or atypical) class of pharmaceuticals, their associations with the sleep EEG were diverse. All antipsychotics share the mechanisms of inhibiting dopamine D2 receptors, while displaying different pharmacology across D1, D3, D4 and D5 receptors, as well as on serotonin receptors, adrenergic receptors, M1, and H1 receptors (Mauri et al., 2014). Such polypharmacy on multiple G protein-coupled receptors of antipsychotics potentially underlies their effects on neurophysiology as some of these receptors play a role in sleep regulation (Eder et al., 2003; Popa et al., 2005). For example, animals with a loss of function of the serotonin receptor gene 5-HT1A showed increased NREM duration (Monaca et al., 2003). More generally, our results underscore the importance of more granular control for medication effects in future studies: one of the most common approaches – using the total antipsychotic dosage equivalent to chlorpromazine – lacked significant associations with key sleep metrics, despite clear effects of individual medications. Other medication classes including sedatives, commonly used as adjunct medications in SCZ, had large effects on sleep macro and micro-architecture, as other have noted (Leong et al., 2022). Sedatives increased both slow and fast spindle density, consistent with the premise of prior literature that examined their role in treating memory deficits in SCZ patients (Mylonas et al., 2020a; Wamsley et al., 2013).

Adjusting for medication use accounted for a substantial proportion of the heterogeneity in sleep metrics, based on the number of metrics with significantly increased variance in SCZ. The interpretation of associations with medication use in the sleep EEG is nonetheless challenging: a given effect could be 1) a purely epiphenomenal side-effect, 2) a marker of therapeutic action, normalizing canonical deficits, or 3) a consequence of non-random prescription of particular medications to particular patients, based on clinical course. Indeed, we observed antipsychotics impacting sleep metrics 1) that, on average, did not differ between patients and controls 2) that had the opposite direction of effect or 3) that had the same direction of effect compared to the SCZ/CTR contrast. That is, associations between certain medications and the sleep EEG may not always be simple confounds *per se*, but instead reflect an individual’s particular deficits, an allostatic response to those deficits, or a personalized response to treatment, as well as the specific properties of the particular treatment on other molecular targets.

Adjustment for medication effects still left a significant portion of the between-patient variability in NREM sleep unexplained however, which presumably reflects either genetic, environmental risks, as well as developmental and dynamic sources of intra-individual variability. There are mixed findings regarding the role of genetics in driving between-patient variability: for example, whereas one study found no relation between genetic risk of SCZ and brain structural heterogeneity (Alnæs et al., 2019), another found a significant association between SCZ polygenic risk and a greater number of brain regions displaying deviations in cortical thickness (Lv et al., 2021).

One possible driver of higher between-individual variability in SCZ is greater reactivity to other exogeneous factors that impact sleep in the general population, including demographics. Indeed, for at least some key metrics patients tended to exhibit greater age-related changes for some key metrics (fast spindle density, total sleep time), although this pattern was not routinely observed for all metrics displaying comparably large case-control differences. Based on an independent prediction model of brain age from the sleep EEG, patients showed profiles of NREM sleep metrics consistent with accelerated ageing. Although the extent to which biological age as predicted from the sleep EEG captures the same phenomena as age predicted from MRI imaging (or other modalities including epigenetics) is unclear, our findings were consistent with a recent meta-analysis that found increased brain ages in patients compared to controls, but no relationship with clinical factors in patients (Constantinides et al., 2023). Also consistent with (Constantinides et al., 2023), there was no association between PAD and CPZ equivalent dose, although we did find a large effect of olanzapine use, with use being associated with advanced ageing. An important caveat is that factors such as olanzapine use were likely not represented in the training data for the age prediction model used here: that is, the NREM EEG signature of olanzapine use may confound the model’s predictions. Future work including longitudinal approaches will be necessary to determine whether this increased inter-individual variability is also reflected in altered intra-individual variability, that is, the temporal stability of the sleep EEG over different timescales, from years to days to seconds.

Our results continue to support the relevance of the distinction between fast and slow spindles: for example, in patients we observed qualitatively different patterns of results for these spindle classes with respect to patterns of age-related decline in spindle density, as well as differential associations with cognitive versus clinical features. We further identified instances where the topographic and temporal context of spindles played a critical role, including the relative increase in SO-coupled occipital fast spindles in patients, potentially reflecting the potentiating effect of SO state on spindle generation, which may become a more critical factor in individuals with otherwise disrupted spindle-generating circuits.

Our study is not without caveats and limitations. Of note, the inpatient context likely impacts the extent to which circadian factors and daily rest/activity rhythms play a role. Although we attempted to document and control for medication effects among patients, it is still challenging to fully account for such effects, and medications used by only a handful of patients were excluded from analyses. Translational research using animal models may help address the impact of acute or chronic administration of these antipsychotics and adjunct medications. In addition, characteristics of drug naïve patients and longitudinal studies may provide evidence of sleep EEG metrics changes before and after medication use. Finally, although previous studies reported night-to-night stability in SCZ with respect to spindle deficits over short timescales (Cox et al., 2017; Mylonas et al., 2020b), our restriction to a single night still limits our ability to disentangle within-patient night-to-night variability (over timescales from days to years) from more “trait-like” between-patient factors.

In summary, the current study offers substantial evidence of robust and replicable alterations in sleep neurophysiology as well as increased variability among SCZ patients. Part of this increased variability may be explained by an acceleration of normal age-related changes in patients. Part – but not all – may be explained by patterns of medication use, that will be important to more directly model in future studies aiming to more precisely link clinical and cognitive outcomes to sleep physiology. Whereas group-level mean differences in NREM neurophysiology have now been unambiguously established, the substantial heterogeneity in sleep architecture among patients, as well as their cognitive and clinical symptoms, points to the need for large, transdiagnostic, demographically diverse, genotypically informative and deeply phenotyped samples to characterize the underlying links between sleep and individual patient characteristics.

## METHODS

### Participants

Data on 301 individuals (175 SCZ and 126 CTR) were collected as part of the Global Research Initiative of Neurophysiology on Schizophrenia (GRINS) study. A portion of these data (N=130, Wave 1) was collected before October 12, 2020 and was used in our prior report (Kozhemiako et al., 2022). Since then, the second wave of data (171 additional individuals, Wave 2) was collected following the same protocol and inclusion/exclusion criteria as described in Kozhemiako et al., 2022. In brief, all participants were aged 18-45 with normal IQ (>70). Patients with schizophrenia (SCZ) were recruited from Wuxi Mental Health Center and diagnosed according to DSM-5. Control subjects, without any mental disorders or family history thereof, were recruited from the local community through advertisements. Additionally, the following exclusion criteria applied to all participants: (1) less than 6 months since electroconvulsive treatment; (2) self-reported sleep disorders or barbiturate use; (3) severe medical conditions like epilepsy or head injury; (4) hearing impairment (above 45 dB at 1000 Hz); and (5) pregnancy or lactation. Informed written consent was given by all participants. The study conformed to the Declaration of Helsinki and was approved by the Harvard TH Chan School of Public Health Office of Human Research Administration (IRB18-0058) as well as the Institutional Review Board of WMHC (WXMHCIRB2018LLKY003).

### Data acquisition

All participants underwent three separate visits: 1) to determine eligibility, 2) clinical assessments and the collection of demographic and other medical information, and 3) an overnight EEG, including an event-related potential (ERP) session in the evening. The ERP paradigms included sensory gating, auditory 40 Hz steady-state response and mismatch negativity. EEG recordings used a customized 64-channel EasyCap and the BrainAmp Standard recorder (manufactured by Brain Products GmbH, Germany) at a sampling rate of 500 Hz.

Diagnoses of SCZ were validated using the Structured Clinical Interview for DSM Disorders (SCID)(Phillips et al., 2009). Control subjects were screened by a psychiatrist to confirm the absence of major mental disorders. Collection of clinical information (the Positive and Negative Syndrome Scale (PANSS) (Kay et al., 1987)) and cognitive assessments (the MATRICS Consensus Cognitive Battery (MCCB)(August et al., 2012) was conducted by a full-time researcher. PANSS and SCID assessments were conducted by trained psychiatrists, within a week of the sleep EEG.

### MST task description

The Finger Tapping Motor Sequence Task (MST) involves quickly and accurately pressing four labeled keys on a computer keyboard with the left hand in a 5-element sequence for 30 seconds (a trial). The MST included training and testing runs and was run twice: overnight session with training run in the evening (12 trials) and testing run (12 trials) in the morning after sleep and morning control session with training (12 trials) and testing (6 trials) runs in the morning with only 10 minutes in between. Different sequences were used for overnight and morning 10-min MST sessions and reward was offered for total correct sequences to motivate performance. The primary measure is the number of correct sequences per trial, reflecting both speed and accuracy. Overnight improvement is calculated as the percentage increase in correct sequences from the best three training trials to the first three test trials the next morning. For consistency with prior literature, the last three training trials were also used to compute percentage improvement but it did not alter the results. Learning rate was computed as an average of the best three training trials over the first trial performance.

### Sleep EEG analysis

An open-source package Luna (http://zzz.bwh.harvard.edu/luna/) developed by us (SMP) was used process the sleep EEG data. EEG channels were re-referenced to linked mastoids, down sampled to 200 Hz and band-pass filtered (0.3-35 Hz). Subsequently, all epochs of a specific manually-scored stage (all primary analyses focus on N2) had outliers removed or interpolated based on the next steps. Firstly, we initiated the process of detecting channels with significant and persistent artifacts. These problematic channels were subsequently interpolated using spherical spline interpolation (Perrin et al., 1989). Specifically, a channel was designated as bad if over 30% of its data epochs deviated by more than 2 standard deviations from the mean of all channels, in relation to any of the three Hjorth parameters: activity, mobility, and complexity, as originally proposed by (Hjorth, 1970). This comparison was made within each epoch across all channels. Secondly, we identified outlier data epochs by comparing them to all other epochs recorded from all EEG channels. This comparison was done using the same Hjorth criteria, but we used a more rigorous threshold of 4 standard deviations. Additionally, any epochs with maximum amplitudes exceeding 500 μV, or those exhibiting flat or clipped signals for more than 10% of the epoch’s duration, were also marked as outliers and subsequently interpolated. On average, 7.6% and 7.8% epochs were removed in SCZ and CTR groups respectively in Wave 2, leaving 386 (43-751) and 400 (92-562) epochs in each group. There was no significant group differences (p > 0.05) in the proportion of epochs removed or the number epochs.

### Spindle, SOs detection and coupling estimation

Spindles were detected using a wavelet method with specific center frequencies of 11 Hz for SS and 15 Hz for FS (targeting approximately +/− 2 Hz). Putative spindles were identified based on exceeding certain thresholds and temporal criteria described in detail in Kozhemiako et.al 2022. Non-spindle bands activity increase was compared to spindle frequency activity to ensure spindles primarily reflect sigma-band activity. For qualified spindles, we estimated density, amplitude, ISA, duration, observed frequency, and chirp.

SOs were identified by detecting zero-crossings in the 0.3-4 Hz bandpass-filtered EEG signals based on specific temporal criteria (zero-crossing leading to negative peak was between 0.3 and 1.5 s long; a zero-crossing leading to positive peak were not longer than 1 s) and adaptive/relative amplitude thresholds (twice the size of the signal mean). For each channel, SO density, negative-peak and peak-to-peak amplitude, duration, and upward slope of the negative peak were computed.

To assess SO/spindle coupling, SO phase at the spindle peak was estimated using a filter-Hilbert method: the circular mean SO phase (angle) and inter-trial phase clustering (magnitude) metric were used to quantify consistency of coupling. SO/spindle overlap was also measured as the proportion of spindles overlapping with a detected SO. To account for differences in spindle and SO density, we used randomized surrogate time series to express magnitude and overlap metrics as Z-scores, relative to the empirical null distribution. SO phase-spindle frequency modulation was assessed by a circular-linear correlation, with SO phase split into 18 20-degree bins and instantaneous spindle frequency averaged across bins.

### Spectral power and functional connectivity

Spectral power was estimated using Welch’s method, averageing 0.5 to 20 Hz power spectra across 4-second segments within each 30-second epoch. To assess connectivity between channels, phase slope index (PSI) values were calculated for all channel pairs within 10-minutes of randomly selected N2 sleep epochs, and normalized based on the SD. Only channel wise net PSI values (the sum of all PSI values for a given channel) are reported here, to reflect whether it was predominantly as a sender or recipient of information from other channels.

### Biological age prediction based on the sleep EEG

We used a modified version of the model described in (Sun et al., 2019). This model used 13 features from the sleep EEG and was trained on over 2,500 individuals aged 18 to 80 from the Massachusetts General Hospital sleep clinic. The revised model features and weights are available as part of the Luna package (https://zzz.bwh.harvard.edu/luna/ref/predict/). The model uses 13 features from the NREM EEG based on two central mastoid-referenced channels (C3 and C4): mean band power (N3 delta &N1 alpha), spectral kurtosis (N2 delta, theta, alpha & sigma; N3 theta), time-domain kurtosis (N2 and N3), band power ratios (N3 delta/theta and delta/alpha), *F_C_*= 13.5 Hz spindle density and number of spindles overlapping a detected SO.

### Statistical analyses: group differences and association analyses

To examine if sleep metrics were different between groups or their association with non-sleep variables were significant, we used linear regression models incorporating age and sex as covariates:

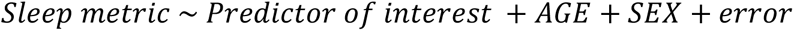

where the predictor of interest was either diagnostic status (SCZ vs CTR) or non-sleep variables such as clinical factors, medication, and cognitive scores. Outlier EEG metric values (> 3 SD from the mean) were removed prior to analysis.

### Accounting for multiple comparisons

To account for comparisons for EEG metrics defined across multiple channels and potentially also multiple frequencies, we utilized a cluster-based permutation using the Freedman-Lane method to correct for nuisance variables (Freedman and Lane, 1983; Winkler et al., 2014), as implemented in Luna. We used a clustering heuristic to identify groups of adjacent predictors and tested significance empirically. Adjacency was determined with respect to spatial location (< 0.5 Euclidian distance of channels) and, for some metrics, also frequency (< 0.5 Hz for PSD, < 1 Hz for PSI). Clusters were defined based on an absolute *t*-score threshold *t*=2. 3000 permutations (based on permuting observed residuals following Freedman-Lane) were used to construct a null distribution to assess statistical significance clusters. Participants were excluded if one or more features were outliers (6 SD for PSD and PSI, 4 SD for all other metrics).

The above approach controls for type I rate at the level of an individual metric, adjusting for the multiple correlated channels (and possibly frequencies) tested. To adjust for testing multiple classes of metric (**Table 2**), we additionally applied FDR correction (alpha=0.05) across all comparisons represented in each matrix of Figure 3 A, B, and Figure 4 A. For multichannel metrics, *t*-scores were presented for the channel with the largest absolute t-score; a metric was considered significant if it both belonged to a significant cluster (*p* < 0.05) and that channel’s FDR-adjusted asymptotic *p*-value was < 0.05.

### Case-control classification

As previously reported (Kozhemiako et al., 2022), a logistic regression model was trained using Wave 1 data to classify SCZ and CTR subjects based on 12 principal components (PCs, see Table 2). After projecting all wave 2 individuals into this PC space, we computed the probability of being a case, based on the previous model. Age and sex effects were regressed out before fitting the model. Model performance was evaluated using the area under the ROC curve (AUC).

### Inter-subject variability analysis

Group differences in the inter-subject variability were tested using Bartlett’s test for homogeneity of variance, with the effects of sex and age regressed out and outlier values (> 3 SD from the mean) removed. To estimate the extent to which clinical, cognitive or medication effects contributed to increased between-individual variability in SCZ, we repeated Bartlett’s tests on residuals also accounting for that class of measure. Specifically, for clinical factors, we controlled for illness duration, total antipsychotics dosage and 5-factor severity scores; for cognitive factors, we controlled for MST overnight and morning test performance, MCCB composite and domain scores; for medication effects, we controlled for binary variables indicating the use of a particular type of medication. (Note: we only considered medication groups if 10 or more patients were using it.). For each sleep metric, we first estimated residuals from a model combining patients and controls with age and sex as predictors, and then further adjusted based on a second model for SCZ individuals only, to remove additional variance associated with clinical, cognitive or medication factors. The residuals then were tested using Bartlett’s test.

### Cluster analysis

In exploratory analyses, SCZ individuals were clustered using dimensionality reduction techniques UMAP and PCA, applied to selected sets of sleep metrics: those with increased variance in SCZ, or those showing SCZ-CTR mean differences at various significance thresholds (p < 0.05, p < 0.01, p < 0.001).

## Acknowledgements

The GRINS study is funded by the Stanley Center for Psychiatric Research and Wuxi Mental Health Center. This work was additionally supported by R01 MH115045-01 and R01MH118298 to JQP; by R03 MH108908, R01 HL146339, R21 HL145492 and R21 MD012738 to SMP; by R01MH092638 and UG3 MH125273 to D.S.M; by K23MH118565 to M.M; by Brain & Behavior Research Foundation Young Investigator and the Zhengxu and Ying He Foundation awards to HH; by the Wuxi City Young Investigator Award to JW.

## Notes

### Competing Interest Statement

The authors have declared no competing interest.

## References

Ahmed, A.O., Strauss, G.P., Buchanan, R.W., Kirkpatrick, B., Carpenter, W.T., 2018. Schizophrenia heterogeneity revisited: Clinical, cognitive, and psychosocial correlates of statistically-derived negative symptoms subgroups. J Psychiatr Res 97, 8–15. 10.1016/j.jpsychires.2017.11.004

Alnæs, D., Kaufmann, T., van der Meer, D., Córdova-Palomera, A., Rokicki, J., Moberget, T., Bettella, F., Agartz, I., Barch, D.M., Bertolino, A., Brandt, C.L., Cervenka, S., Djurovic, S., Doan, N.T., Eisenacher, S., Fatouros-Bergman, H., Flyckt, L., Di Giorgio, A., Haatveit, B., Jönsson, E.G., Kirsch, P., Lund, M.J., Meyer-Lindenberg, A., Pergola, G., Schwarz, E., Smeland, O.B., Quarto, T., Zink, M., Andreassen, O.A., Westlye, L.T., for the Karolinska Schizophrenia Project Consortium, 2019. Brain Heterogeneity in Schizophrenia and Its Association With Polygenic Risk. JAMA Psychiatry 76, 739–748. 10.1001/jamapsychiatry.2019.0257

Arai, Y., Sasayama, D., Kuraishi, K., Murata, S., Usuda, N., Tsuchida, M., Nakajima, Y., Washizuka, S., 2023. Analysis of the effect of brexpiprazole on sleep architecture in patients with schizophrenia: A preliminary study. Neuropsychopharmacology Reports 43, 112–119. 10.1002/npr2.12317

Au, C.H., Harvey, C.-J., 2020. Systematic review: the relationship between sleep spindle activity with cognitive functions, positive and negative symptoms in psychosis. Sleep Medicine: X 2, 100025. 10.1016/j.sleepx.2020.100025

August, S.M., Kiwanuka, J.N., McMahon, R.P., Gold, J.M., 2012. The MATRICS Consensus Cognitive Battery (MCCB): Clinical and Cognitive Correlates. Schizophr Res 134, 76–82. 10.1016/j.schres.2011.10.015

Baandrup, L., Christensen, J.A.E., Fagerlund, B., Jennum, P., 2019. Investigation of sleep spindle activity and morphology as predictors of neurocognitive functioning in medicated patients with schizophrenia. Journal of Sleep Research 28, e12672. 10.1111/jsr.12672

Bagautdinova, J., Mayeli, A., Wilson, J.D., Donati, F.L., Colacot, R.M., Meyer, N., Fusar-Poli, P., Ferrarelli, F., 2023. Sleep Abnormalities in Different Clinical Stages of Psychosis: A Systematic Review and Meta-analysis. JAMA Psychiatry 80, 202–210. 10.1001/jamapsychiatry.2022.4599

Brugger, S.P., Howes, O.D., 2017. Heterogeneity and Homogeneity of Regional Brain Structure in Schizophrenia: A Meta-analysis. JAMA Psychiatry 74, 1104–1111. 10.1001/jamapsychiatry.2017.2663

Carruthers, S.P., Brunetti, G., Rossell, S.L., 2021. Sleep disturbances and cognitive impairment in schizophrenia spectrum disorders: a systematic review and narrative synthesis. Sleep Medicine 84, 8–19. 10.1016/j.sleep.2021.05.011

Chan, M.-S., Chung, K.-F., Yung, K.-P., Yeung, W.-F., 2017. Sleep in schizophrenia: A systematic review and meta-analysis of polysomnographic findings in case-control studies. Sleep Medicine Reviews 32, 69–84. 10.1016/j.smrv.2016.03.001

Constantinides, C., Han, L.K.M., Alloza, C., Antonucci, L.A., Arango, C., Ayesa-Arriola, R., Banaj, N., Bertolino, A., Borgwardt, S., Bruggemann, J., Bustillo, J., Bykhovski, O., Calhoun, V., Carr, V., Catts, S., Chung, Y.-C., Crespo-Facorro, B., Díaz-Caneja, C.M., Donohoe, G., Plessis, S.D., Edmond, J., Ehrlich, S., Emsley, R., Eyler, L.T., Fuentes-Claramonte, P., Georgiadis, F., Green, M., Guerrero-Pedraza, A., Ha, M., Hahn, T., Henskens, F.A., Holleran, L., Homan, S., Homan, P., Jahanshad, N., Janssen, J., Ji, E., Kaiser, S., Kaleda, V., Kim, M., Kim, W.-S., Kirschner, M., Kochunov, P., Kwak, Y.B., Kwon, J.S., Lebedeva, I., Liu, J., Mitchie, P., Michielse, S., Mothersill, D., Mowry, B., de la Foz, V.O.-G., Pantelis, C., Pergola, G., Piras, F., Pomarol-Clotet, E., Preda, A., Quidé, Y., Rasser, P.E., Rootes-Murdy, K., Salvador, R., Sangiuliano, M., Sarró, S., Schall, U., Schmidt, A., Scott, R.J., Selvaggi, P., Sim, K., Skoch, A., Spalletta, G., Spaniel, F., Thomopoulos, S.I., Tomecek, D., Tomyshev, A.S., Tordesillas-Gutiérrez, D., van Amelsvoort, T., Vázquez-Bourgon, J., Vecchio, D., Voineskos, A., Weickert, C.S., Weickert, T., Thompson, P.M., Schmaal, L., van Erp, T.G.M., Turner, J., Cole, J.H., ENIGMA Schizophrenia Consortium, Dima, D., Walton, E., 2023. Brain ageing in schizophrenia: evidence from 26 international cohorts via the ENIGMA Schizophrenia consortium. Mol Psychiatry 28, 1201–1209. 10.1038/s41380-022-01897-w

Cox, R., Schapiro, A.C., Manoach, D.S., Stickgold, R., 2017. Individual Differences in Frequency and Topography of Slow and Fast Sleep Spindles. Front. Hum. Neurosci. 11. 10.3389/fnhum.2017.00433

Demanuele, C., Bartsch, U., Baran, B., Khan, S., Vangel, M.G., Cox, R., Hämäläinen, M., Jones, M.W., Stickgold, R., Manoach, D.S., 2017. Coordination of Slow Waves With Sleep Spindles Predicts Sleep-Dependent Memory Consolidation in Schizophrenia. Sleep 40. 10.1093/sleep/zsw013

Demirlek, C., Bora, E., 2023. Sleep-dependent memory consolidation in schizophrenia: A systematic review and meta-analysis. Schizophrenia Research 254, 146–154. 10.1016/j.schres.2023.02.028

Demjaha, A., Lappin, J.M., Stahl, D., Patel, M.X., MacCabe, J.H., Howes, O.D., Heslin, M., Reininghaus, U.A., Donoghue, K., Lomas, B., Charalambides, M., Onyejiaka, A., Fearon, P., Jones, P., Doody, G., Morgan, C., Dazzan, P., Murray, R.M., 2017. Antipsychotic treatment resistance in first-episode psychosis: prevalence, subtypes and predictors. Psychol Med 47, 1981–1989. 10.1017/S0033291717000435

Eder, D.N., Zdravkovic, M., Wildschiødtz, G., 2003. Selective alterations of the first NREM sleep cycle in humans by a dopamine D1 receptor antagonist (NNC-687). J Psychiatr Res 37, 305–312. 10.1016/s0022-3956(03)00007-4

Erickson, M.A., Ruffle, A., Gold, J.M., 2016. A meta-analysis of mismatch negativity in schizophrenia: from clinical risk to disease specificity and progression. Biol Psychiatry 79, 980–987. 10.1016/j.biopsych.2015.08.025

Ferrarelli, F., 2023. Sleep spindles as neurophysiological biomarkers of schizophrenia. Eur J Neurosci. 10.1111/ejn.16178

Freedman, D., Lane, D., 1983. A Nonstochastic Interpretation of Reported Significance Levels. Journal of Business & Economic Statistics 1, 292–298. 10.1080/07350015.1983.10509354

Freedman, R., Olsen-Dufour, A.M., Olincy, A., 2020. P50 inhibitory sensory gating in schizophrenia: analysis of recent studies. Schizophrenia Research 218, 93–98. 10.1016/j.schres.2020.02.003

Göder, R., Fritzer, G., Gottwald, B., Lippmann, B., Seeck-Hirschner, M., Serafin, I., Aldenhoff, J.B., 2008. Effects of olanzapine on slow wave sleep, sleep spindles and sleep-related memory consolidation in schizophrenia. Pharmacopsychiatry 41, 92–99. 10.1055/s-2007-1004592

Gopal, S., Miller, R.L., Michael, A., Adali, T., Cetin, M., Rachakonda, S., Bustillo, J.R., Cahill, N., Baum, S.A., Calhoun, V.D., 2016. Spatial Variance in Resting fMRI Networks of Schizophrenia Patients: An Independent Vector Analysis. Schizophrenia Bulletin 42, 152–160. 10.1093/schbul/sbv085

Hjorth, B., 1970. EEG analysis based on time domain properties. Electroencephalography and Clinical Neurophysiology 29, 306–310. 10.1016/0013-4694(70)90143-4

Hjorthøj, C., Stürup, A.E., McGrath, J.J., Nordentoft, M., 2017. Years of potential life lost and life expectancy in schizophrenia: a systematic review and meta-analysis. Lancet Psychiatry 4, 295–301. 10.1016/S2215-0366(17)30078-0

International Schizophrenia Consortium, Purcell, S.M., Wray, N.R., Stone, J.L., Visscher, P.M., O’Donovan, M.C., Sullivan, P.F., Sklar, P., 2009. Common polygenic variation contributes to risk of schizophrenia and bipolar disorder. Nature 460, 748–752. 10.1038/nature08185

Kaskie, R.E., Ferrarelli, F., 2020. Sleep disturbances in schizophrenia: what we know, what still needs to be done. Current Opinion in Psychology, Sleep & Psychopathology 34, 68–71. 10.1016/j.copsyc.2019.09.011

Kaufmann, T., van der Meer, D., Doan, N.T., Schwarz, E., Lund, M.J., Agartz, I., Alnæs, D., Barch, D.M., Baur-Streubel, R., Bertolino, A., Bettella, F., Beyer, M.K., Bøen, E., Borgwardt, S., Brandt, C.L., Buitelaar, J., Celius, E.G., Cervenka, S., Conzelmann, A., Córdova-Palomera, A., Dale, A.M., de Quervain, D.J.F., Di Carlo, P., Djurovic, S., Dørum, E.S., Eisenacher, S., Elvsåshagen, T., Espeseth, T., Fatouros-Bergman, H., Flyckt, L., Franke, B., Frei, O., Haatveit, B., Håberg, A.K., Harbo, H.F., Hartman, C.A., Heslenfeld, D., Hoekstra, P.J., Høgestøl, E.A., Jernigan, T.L., Jonassen, R., Jönsson, E.G., Kirsch, P., Kłoszewska, I., Kolskår, K.K., Landrø, N.I., Le Hellard, S., Lesch, K.-P., Lovestone, S., Lundervold, A., Lundervold, A.J., Maglanoc, L.A., Malt, U.F., Mecocci, P., Melle, I., Meyer-Lindenberg, A., Moberget, T., Norbom, L.B., Nordvik, J.E., Nyberg, L., Oosterlaan, J., Papalino, M., Papassotiropoulos, A., Pauli, P., Pergola, G., Persson, K., Richard, G., Rokicki, J., Sanders, A.-M., Selbæk, G., Shadrin, A.A., Smeland, O.B., Soininen, H., Sowa, P., Steen, V.M., Tsolaki, M., Ulrichsen, K.M., Vellas, B., Wang, L., Westman, E., Ziegler, G.C., Zink, M., Andreassen, O.A., Westlye, L.T., 2019. Common brain disorders are associated with heritable patterns of apparent aging of the brain. Nat Neurosci 22, 1617–1623. 10.1038/s41593-019-0471-7

Kay, S., Fiszbein, A., Opler, L., 1987. The Positive and Negative Syndrome Scale (PANSS) for schizophrenia. Schizophrenia bulletin 13, 261–76. 10.1093/schbul/13.2.261

Kluge, M., Schacht, A., Himmerich, H., Rummel-Kluge, C., Wehmeier, P.M., Dalal, M., Hinze-Selch, D., Kraus, T., Dittmann, R.W., Pollmächer, T., Schuld, A., 2014. Olanzapine and clozapine differently affect sleep in patients with schizophrenia: results from a double-blind, polysomnographic study and review of the literature. Schizophr Res 152, 255–260. 10.1016/j.schres.2013.11.009

Kozhemiako, N., Buckley, A.W., Chervin, R.D., Redline, S., Purcell, S.M., 2023. Mapping Neurodevelopment with Sleep Macro-and Micro-Architecture Across Multiple Pediatric Populations. NeuroImage: Clinical 103552. 10.1016/j.nicl.2023.103552

Kozhemiako, N., Wang, J., Jiang, C., Wang, L.A., Gai, G., Zou, K., Wang, Z., Yu, X., Zhou, L., Li, S., Guo, Z., Law, R., Coleman, J., Mylonas, D., Shen, L., Wang, G., Tan, S., Qin, S., Huang, H., Murphy, M., Stickgold, R., Manoach, D., Zhou, Z., Zhu, W., Hal, M.-H., Purcell, S.M., Pan, J.Q., 2022. Non-rapid eye movement sleep and wake neurophysiology in schizophrenia. eLife 11, e76211. 10.7554/eLife.76211

Krystal, A.D., Goforth, H.W., Roth, T., 2008. Effects of antipsychotic medications on sleep in schizophrenia: International Clinical Psychopharmacology 23, 150–160. 10.1097/YIC.0b013e3282f39703

Lai, M., Hegde, R., Kelly, S., Bannai, D., Lizano, P., Stickgold, R., Manoach, D.S., Keshavan, M., 2022. Investigating sleep spindle density and schizophrenia: A meta-analysis. Psychiatry Research 307, 114265. 10.1016/j.psychres.2021.114265

Leong, C.W.Y., Leow, J.W.S., Grunstein, R.R., Naismith, S.L., Teh, J.Z., D’Rozario, A.L., Saini, B., 2022. A systematic scoping review of the effects of central nervous system active drugs on sleep spindles and sleep-dependent memory consolidation. Sleep Medicine Reviews 62, 101605. 10.1016/j.smrv.2022.101605

Lv, J., Di Biase, M., Cash, R.F.H., Cocchi, L., Cropley, V.L., Klauser, P., Tian, Y., Bayer, J., Schmaal, L., Cetin-Karayumak, S., Rathi, Y., Pasternak, O., Bousman, C., Pantelis, C., Calamante, F., Zalesky, A., 2021. Individual deviations from normative models of brain structure in a large cross-sectional schizophrenia cohort. Mol Psychiatry 26, 3512–3523. 10.1038/s41380-020-00882-5

Manoach, D.S., Cain, M.S., Vangel, M.G., Khurana, A., Goff, D.C., Stickgold, R., 2004. A failure of sleep-dependent procedural learning in chronic, medicated schizophrenia. Biological Psychiatry 56, 951–956. 10.1016/j.biopsych.2004.09.012

Manoach, D.S., Demanuele, C., Wamsley, E.J., Vangel, M., Montrose, D.M., Miewald, J., Kupfer, D., Buysse, D., Stickgold, R., Keshavan, M.S., 2014. Sleep spindle deficits in antipsychotic-naïve early course schizophrenia and in non-psychotic first-degree relatives. Front. Hum. Neurosci. 8. 10.3389/fnhum.2014.00762

Manoach, D.S., Stickgold, R., 2019. Abnormal Sleep Spindles, Memory Consolidation, and Schizophrenia. Annual Review of Clinical Psychology 15, 451–479. 10.1146/annurev-clinpsy-050718-095754

Manoach, D.S., Thakkar, K.N., Stroynowski, E., Ely, A., McKinley, S.K., Wamsley, E., Djonlagic, I., Vangel, M.G., Goff, D.C., Stickgold, R., 2010. Reduced overnight consolidation of procedural learning in chronic medicated schizophrenia is related to specific sleep stages. Journal of Psychiatric Research 44, 112–120. 10.1016/j.jpsychires.2009.06.011

Mauri, M.C., Paletta, S., Maffini, M., Colasanti, A., Dragogna, F., Di Pace, C., Altamura, A.C., 2014. Clinical pharmacology of atypical antipsychotics: an update. EXCLI J 13, 1163–1191.

Monaca, C., Boutrel, B., Hen, R., Hamon, M., Adrien, J., 2003. 5-HT1A/1B Receptor-Mediated Effects of the Selective Serotonin Reuptake Inhibitor, Citalopram, on Sleep: Studies in 5-HT1A and 5-HT1B Knockout Mice. Neuropsychopharmacol 28, 850–856. 10.1038/sj.npp.1300109

Mylonas, D., Baran, B., Demanuele, C., Cox, R., Vuper, T.C., Seicol, B.J., Fowler, R.A., Correll, D., Parr, E., Callahan, C.E., Morgan, A., Henderson, D., Vangel, M., Stickgold, R., Manoach, D.S., 2020a. The effects of eszopiclone on sleep spindles and memory consolidation in schizophrenia: a randomized clinical trial. Neuropsychopharmacol. 45, 2189–2197. 10.1038/s41386-020-00833-2

Mylonas, D., Tocci, C., Coon, W.G., Baran, B., Kohnke, E.J., Zhu, L., Vangel, M.G., Stickgold, R., Manoach, D.S., 2020b. Naps reliably estimate nocturnal sleep spindle density in health and schizophrenia. J Sleep Res 29, e12968. 10.1111/jsr.12968

Perrin, F., Pernier, J., Bertrand, O., Echallier, J.F., 1989. Spherical splines for scalp potential and current density mapping. Electroencephalogr Clin Neurophysiol 72, 184–187. 10.1016/0013-4694(89)90180-6

Phillips, M.R., Zhang, J., Shi, Q., Song, Z., Ding, Z., Pang, S., Li, X., Zhang, Y., Wang, Z., 2009. Prevalence, treatment, and associated disability of mental disorders in four provinces in China during 2001-05: an epidemiological survey. Lancet 373, 2041–2053. 10.1016/S0140-6736(09)60660-7

Popa, D., Léna, C., Fabre, V., Prenat, C., Gingrich, J., Escourrou, P., Hamon, M., Adrien, J., 2005. Contribution of 5-HT2 receptor subtypes to sleep-wakefulness and respiratory control, and functional adaptations in knock-out mice lacking 5-HT2A receptors. J Neurosci 25, 11231–11238. 10.1523/JNEUROSCI.1724-05.2005

Purcell, S.M., Manoach, D.S., Demanuele, C., Cade, B.E., Mariani, S., Cox, R., Panagiotaropoulou, G., Saxena, R., Pan, J.Q., Smoller, J.W., Redline, S., Stickgold, R., 2017. Characterizing sleep spindles in 11,630 individuals from the National Sleep Research Resource. Nature Communications 8, 15930. 10.1038/ncomms15930

Seaton, B.E., Goldstein, G., Allen, D.N., 2001. Sources of Heterogeneity in Schizophrenia: The Role of Neuropsychological Functioning. Neuropsychol Rev 11, 45–67. 10.1023/A:1009013718684

Stanley, S., Balakrishnan, S., Ilangovan, S., 2017. Psychological distress, perceived burden and quality of life in caregivers of persons with schizophrenia. Journal of Mental Health 26, 134–141. 10.1080/09638237.2016.1276537

Sun, H., Paixao, L., Oliva, J.T., Goparaju, B., Carvalho, D.Z., van Leeuwen, K.G., Akeju, O., Thomas, R.J., Cash, S.S., Bianchi, M.T., Westover, M.B., 2019. Brain age from the electroencephalogram of sleep. Neurobiology of Aging 74, 112–120. 10.1016/j.neurobiolaging.2018.10.016

Thuné, H., Recasens, M., Uhlhaas, P.J., 2016. The 40-Hz Auditory Steady-State Response in Patients With Schizophrenia: A Meta-analysis. JAMA Psychiatry 73, 1145. 10.1001/jamapsychiatry.2016.2619

Trubetskoy, V., Panagiotaropoulou, G., Awasthi, S., Braun, A., Kraft, J., Skarabis, N., Walter, H., Ripke, S., Pardiñas, A.F., Dennison, C.A., Hall, L.S., Harwood, J.C., Richards, A.L., Legge, S.E., Lynham, A., Williams, N.M., Bray, N.J., Escott-Price, V., Kirov, G., Holmans, P.A., Pocklington, A.J., Owen, M.J., Walters, J.T.R., O’Donovan, M.C., Qi, T., Sidorenko, J., Wu, Y., Zeng, J., Gratten, J., Visscher, P.M., Yang, J., Wray, N.R., Qi, T., Yang, J., Bigdeli, T.B., Fanous, A.H., Bryois, J., Bergen, S.E., Kähler, A.K., Magnusson, P.K.E., Hultman, C.M., Sullivan, P.F., Chen, C.-Y., Atkinson, E.G., Goldstein, J.I., Howrigan, D.P., Martin, A.R., Daly, M.J., Huang, H., Neale, B.M., Ripke, S., Ge, T., Lam, M., Ge, T., Atkinson, E.G., Belliveau, R.A., Chambert, K.D., Genovese, G., Lee, P.H., Martin, A.R., Pietiläinen, O., McCarroll, S.A., Moran, J.L., Smoller, J.W., Brown, T.C., Feng, G., Hyman, S.E., Sheng, M., Hyman, S.E., Huang, H., Neale, B.M., Lam, M., Chong, S.A., Subramaniam, M., Lam, M., Lencz, T., Malhotra, A.K., Watanabe, K., Frei, O., Agartz, I., Athanasiu, L., Melle, I., Andreassen, O.A., Frei, O., Athanasiu, L., Melle, I., Steen, N.E., Andreassen, O.A., Frei, O., Ge, T., DeLisi, L.E., Mesholam-Gately, R.I., Seidman, L.J., Koopmans, F., Magnusson, S., Stefánsson, H., Stefansson, K., Grove, J., Agerbo, E., Als, T.D., Bybjerg-Grauholm, J., Demontis, D., Hougaard, D.M., Mors, O., Mortensen, P.B., Nordentoft, M., Børglum, A.D., Grove, J., Als, T.D., Demontis, D., Mattheisen, M., Børglum, A.D., Grove, J., Als, T.D., Demontis, D., Børglum, A.D., Kim, M., Gandal, M.J., Li, Z., Shi, Y., Shi, Y., Li, Z., Zhou, W., Qin, S., Shi, Y., Shi, Y., Voloudakis, G., Zhang, W., Roussos, Panos, Zhang, W., Adams, M., McIntosh, A., Agartz, I., Agartz, I., Söderman, E., Jönsson, E.G., McGrath, J.J., Al Eissa, M., Bass, N.J., Fiorentino, A., O’Brien, N.L., Pimm, J., Sharp, S.I., McQuillin, A., Albus, M., Alexander, M., Alizadeh, B.Z., Bruggeman, R., Alizadeh, B.Z., Alptekin, K., Alptekin, K., Amin, F., Arolt, V., Lencer, R., Rothermundt, M., Baune, B.T., Arrojo, M., Azevedo, M.H., Bacanu, S.A., Webb, B.T., Wormley, B.K., Riley, B.P., Kendler, K.S., Begemann, M., Mitjans, M., Steixner-Kumar, A.A., Ehrenreich, H., Bene, J., Benyamin, B., Benyamin, B., Benyamin, B., Blasi, G., Rampino, A., Torretta, S., Bertolino, A., Bobes, J., Bobes, J., Bobes, J., Bonassi, S., Bressan, R.A., Gadelha, A., Noto, C., Bressan, R.A., Gadelha, A., Noto, C., Ota, V.K., Santoro, M.L., Belangero, S.I., Bromet, E.J., Bruggeman, R., Buckley, P.F., Buckner, R.L., Bybjerg-Grauholm, J., Hougaard, D.M., Cahn, W., Kahn, R.S., Cahn, W., Cairns, M.J., Scott, R.J., Tooney, P.A., Cairns, M.J., Schall, U., Scott, R.J., Tooney, P.A., Cairns, M.J., Tooney, P.A., Calkins, M.E., Gur, R.E., Gur, R.C., Turetsky, B.I., Carr, V.J., Carr, V.J., Carr, V.J., Castle, D., Harvey, C., Castle, D., Catts, S.V., Catts, S.V., Chan, R.C.K., Chan, R.C.K., Chaumette, B., Kebir, O., Krebs, M.-O., Chaumette, B., Cheng, W., Cheung, E.F.C., Chong, S.A., Subramaniam, M., Cohen, D., Consoli, A., Giannitelli, M., Laurent-Levinson, C., Cohen, D., Consoli, A., Giannitelli, M., Laurent-Levinson, C., Cohen, D., Cordeiro, Q., Costas, J., Curtis, C., Quattrone, D., Breen, G., Collier, D.A., Di Forti, M., Vassos, E., Curtis, C., Mondelli, V., Quattrone, D., van Amelsvoort, T., Di Forti, M., Murray, R.M., Vassos, E., van Amelsvoort, T., Davidson, M., Davis, K.L., Haroutunian, V., Malaspina, D., Reichenberg, A., Siever, L.J., Silverman, J.M., Buxbaum, J.D., Kahn, R.S., de Haan, L., de Haan, L., de Haan, L., de Haan, L., Degenhardt, F., Forstner, A., Nöthen, M.M., DeLisi, L.E., Dickerson, F., Dikeos, D., Papadimitriou, G.N., Dinan, T., Dinan, T., Djurovic, S., Djurovic, S., Duan, J., Gejman, P.V., Sanders, A.R., Duan, J., Gejman, P.V., Sanders, A.R., Ducci, G., Dudbridge, F., Eriksson, J.G., Eriksson, J.G., Eriksson, J.G., Fañanás, L., Fañanás, L., Peñas, J.G., González-Pinto, A., Molto, M.D., Moreno, C., Parellada, M., Sanjuan, J., Crepo-Facorro, B., Mata, I., Arango, C., Faraone, S.V., Forstner, A., Frank, J., Streit, F., Witt, S.H., Rietschel, M., Freimer, N.B., Ophoff, R.A., Freimer, N.B., Fromer, M., Stahl, E.A., Frustaci, A., Gershon, E.S., Giegling, I., Hartmann, A.M., Konte, B., Rujescu, D., Giusti-Rodríguez, P., Szatkiewicz, J.P., Sullivan, P.F., Godard, S., González Peñas, J., Moreno, C., Parellada, M., Arango, C., González-Pinto, A., Gopal, S., Savitz, A., Li, Q.S., Gratten, J., Green, M.F., Nuechterlein, K.H., Sugar, C.A., Green, M.F., Greenwood, T.A., Light, G.A., Swerdlow, N.R., Braff, D., Guillin, O., Campion, D., Guillin, O., Campion, D., Guillin, O., Gülöksüz, S., Luykx, J.J., Rutten, B.P.F., van Amelsvoort, T., van Winkel, R., van Amelsvoort, T., van Winkel, R., Gülöksüz, S., Gutiérrez, B., Hahn, E., Hakonarson, H., Pellegrino, R., Haroutunian, V., Haroutunian, V., Harvey, C., Pantelis, C., Hayward, C., Henskens, F.A., Kelly, B.J., Herms, S., Hoffmann, P., Howrigan, D.P., Fromer, M., Daly, M.J., Ikeda, M., Iwata, N., Iyegbe, C., van Os, J., Joa, I., Julià, A., Marsal, S., Kam-Thong, T., Rautanen, A., Kamatani, Y., Kamatani, Y., Karachanak-Yankova, S., Toncheva, D., Karachanak-Yankova, S., Keller, M.C., Khrunin, A., Limborska, S., Slominsky, P., Kim, S.-W., Klovins, J., Nikitina-Zake, L., Kondratiev, N., Golimbet, V., Kraft, J., Kubo, M., Kučinskas, V., Kučinskiene, Z.A., Kusumawardhani, A., Kuzelova-Ptackova, H., Landi, S., Lazzeroni, L.C., Levinson, D.F., Lazzeroni, L.C., Lee, P.H., Petryshen, T.L., Smoller, J.W., Lehrer, D.S., Lerer, B., Li, Miaoxin, Lieberman, J., Stroup, T.S., Light, G.A., Braff, D., Liu, C.-M., Liu, C.-M., Hwu, H.-G., Lönnqvist, J., Lönnqvist, J., Loughland, C.M., Lubinski, J., Luykx, J.J., Bakker, S., Kahn, R., Luykx, J.J., Luykx, J.J., Macek, M., Mackinnon, A., Mackinnon, A., Maher, B.S., Maier, W., Malaspina, D., Atbaşoğlu, E.C., Mallet, J., Marder, S.R., Martin, A.R., Huang, H., Martorell, L., Muntané, G., Vilella, E., Mattheisen, M., Meier, S., Mattheisen, M., Mattheisen, M., Schulze, T.G., McCarley, R.W., McDonald, C., Donohoe, G., Morris, D.W., McGrath, J.J., Periyasamy, S., Mowry, B.J., Wray, N.R., McGrath, J.J., Medeiros, H., Sobell, J.L., Medeiros, H., Meier, S., Melegh, B., Mesholam-Gately, R.I., Seidman, L.J., Metspalu, A., Milani, L., Esko, T., Michie, P.T., Milanova, V., Molden, E., Molden, E., Molina, E., Molto, M.D., Molto, M.D., Sanjuan, J., Mondelli, V., Morley, C.P., Muntané, G., Murphy, K.C., Myin-Germeys, I., Nenadić, I., Nenadić, I., Nestadt, G., Pulver, A.E., O’Neill, F.A., Oh, S.-Y., Oh, S.-Y., Olincy, A., Freedman, R., Ota, V.K., Santoro, M.L., Belangero, S.I., Pantelis, C., Pantelis, C., Baune, B.T., Pantelis, C., Baune, B.T., Paunio, T., Paunio, T., Periyasamy, S., Mowry, B.J., Perkins, D.O., Sullivan, P.F., Pfuhlmann, B., Pietiläinen, O., Hyman, S.E., Pietiläinen, O., Benner, C., Pirinen, M., Palotie, A., Daly, M.J., Porteous, D., Powell, J., Quattrone, D., Di Forti, M., Quested, D., Quested, D., Radant, A.D., Tsuang, D.W., Radant, A.D., Tsuang, D.W., Rapaport, M.H., Roe, C., Liu, C., Roffman, J.L., Moran, J.L., Roth, J., Gawlik, M., Saker-Delye, S., Salomaa, V., Suvisaari, J., Sanjuan, J., Schall, U., Scott, R.J., Shi, J., Siever, L.J., Silverman, J.M., Sigurdsson, E., Sigurdsson, E., Sim, K., Sim, K., Sim, K., So, H.-C., So, H.-C., Stain, H.J., Stain, H.J., Steen, N.E., Jönsson, E.G., Stögmann, E., Zimprich, F., Stone, W.S., Stone, W.S., Straub, R.E., Hyde, T., Jaffe, A., Weinberger, D.R., Strengman, E., Sugar, C.A., Svrakic, D.M., Cloninger, C.R., Ta, T.M.T., Ta, T.M.T., Takahashi, A., Terao, C., Thibaut, F., Thibaut, F., Toncheva, D., Tosato, S., Tura, G.B., Üçok, A., Vaaler, A., Vaaler, A., van Winkel, R., Veijola, J., van Winkel, R., Veijola, J., Waddington, J., Waterreus, A., Morgan, V.A., Waterreus, A., Jablensky, A.V., Morgan, V.A., Weiser, M., Wu, J.Q., Xu, Z., Yolken, R., Zai, C.C., Kennedy, J.L., Zai, C.C., Kennedy, J.L., Zhu, F., Zhu, F., Atbaşoğlu, E.C., Saka, M.C., Ayub, M., Black, D.W., Buccola, N.G., Byerley, W.F., Chen, W.J., Chen, W.J., Crespo-Facorro, B., Crespo-Facorro, B., Galletly, C., Galletly, C., Galletly, C., Gennarelli, M., Gennarelli, M., Hwu, H.-G., Mors, O., Müller-Myhsok, B., Müller-Myhsok, B., Müller-Myhsok, B., Neil, A.L., Nordentoft, M., Pato, M.T., Pato, C.N., Pato, M.T., Pato, C.N., Pirinen, M., Pirinen, M., Schulze, T.G., Schulze, T.G., Schulze, T.G., Stahl, E.A., Esko, T., Stahl, E.A., Wang, S.-H., Xu, S., Xu, S., Xu, S., Adolfsson, R., Bramon, E., Cervilla, J.A., Cichon, S., Cichon, S., Cichon, S., Collier, D.A., Collier, D.A., Corvin, A., Gill, M., Curtis, D., Curtis, D., Domenici, E., Escott-Price, V., Fanous, A.H., Fanous, A.H., Gareeva, A., Khusnutdinova, E., Gareeva, A., Gareeva, A., Khusnutdinova, E., Gareeva, A., Glatt, S.J., Hong, K.S., Knowles, J.A., Knowles, J.A., Lee, J., Lee, J., Lencz, T., Malhotra, A.K., Lencz, T., Malhotra, A.K., Liu, J., Liu, J., Malhotra, D., Menezes, P.R., Nimgaonkar, V., Ophoff, R.A., Ophoff, R.A., Paciga, S.A., Palotie, A., Palotie, A., Ripke, S., Qin, S., Rivera, M., Rivera, M., Schwab, S.G., Schwab, S.G., Serretti, A., Sham, P.C., Sham, P.C., Sham, P.C., Sham, P.C., Sham, P.C., Sham, P.C., Clair, D.S., Tsuang, M.T., Tsuang, M.T., van Os, J., Vawter, M.P., Werge, T., Werge, T., Clair, D.S., Werge, T., Werge, T., van Os, J., Wildenauer, Dieter B., van Os, J., Yu, X., Yue, W., Yu, X., Yue, W., Yue, W., Roussos, Panos, Vassos, E., Verhage, M., Koopmans, F., Sahasrabudhe, D., Toonen, R.F., Verhage, M., Verhage, M., Posthuma, D., Yang, J., Dai, N., Wenwen, Q., Wildenauer, D. B., Dai, N., Wenwen, Q., Wildenauer, D. B., Agiananda, F., Amir, N., Antoni, R., Arsianti, T., Asmarahadi, A., Diatri, H., Djatmiko, P., Irmansyah, I., Khalimah, S., Kusumadewi, I., Kusumaningrum, P., Lukman, P.R., Nasrun, M.W., Safyuni, N.S., Prasetyawan, P., Semen, G., Siste, K., Tobing, H., Widiasih, N., Wiguna, T., Wulandari, D., Evalina, N., Hananto, A.J., Ismoyo, J.H., Marini, T.M., Henuhili, S., Reza, M., Yusnadewi, S., Abyzov, A., Akbarian, S., van Bakel, H., Breen, M., Charney, A., Dracheva, S., Girdhar, K., Hoffman, G., Jiang, Y., Pinto, D., Purcell, S., Roussos, Panagiotis, Wiseman, J., Ashley-Koch, A., Crawford, G., Reddy, T., Brown, M., Grennan, K., Bryois, J., Carlyle, B., Emani, P., Galeev, T., Gerstein, M., Gu, M., Guerra, B., Gursoy, G., Kitchen, R., Lee, D., Li, Mingfeng, Liu, S., Navarro, F., Pan, X., Pochareddy, S., Rozowsky, J., Sestan, N., Sethi, A., Shi, X., Szekely, A., Wang, D., Warrell, J., Weissman, S., Wu, F., Xu, X., Coetzee, G., Farnham, P., Lay, F., Rhie, S., Witt, H., Wood, S., Yao, L., Gandal, M., Polioudakis, D., Swarup, V., Won, H., Giase, G., Jiang, S., Kefi, A., Shieh, A., Goes, F., Zandi, P., Kim, Y., Knowles, J.A., Mattei, E., Purcaro, M., Pratt, H., Peters, M.A., Sanders, S., Weng, Z., White, K., Arranz, M.J., Bramon, E., Iyegbe, C., Lewis, C., Lin, K., Murray, R.M., Powell, J., Walshe, M., Arranz, M.J., Bender, S., Bender, S., Weisbrod, M., Bramon, E., Crepo-Facorro, B., Mata, I., Hall, J., Lawrie, S., McIntosh, A., Linszen, D.H., Ophoff, R.A., van Os, J., van Os, J., Rujescu, D., Rujescu, D., Achsel, T., Bagni, C., Andres-Alonso, M., Kreutz, M.R., Bayés, À., Biederer, T., Brose, N., Chua, J.J.E., Coba, M.P., Cornelisse, L.N., van Weering, J.R.T., de Jong, A.P.H., MacGillavry, H.D., de Juan-Sanz, J., Dieterich, D.C., Pielot, R., Smalla, K.-H., Dieterich, D.C., Gundelfinger, E.D., Pielot, R., Smalla, K.-H., Feng, G., Goldschmidt, H.L., Huganir, R.L., Hoogenraad, C., Hyman, S.E., Imig, C., Jahn, R., Jung, H., Kim, E., Kaeser, P.S., Lipstein, N., Malenka, R., McPherson, P.S., O’Connor, V., Ryan, T.A., Sala, C., Verpelli, C., Smit, A.B., Südhof, T.C., Thomas, P.D., 2022. Mapping genomic loci implicates genes and synaptic biology in schizophrenia. Nature 604, 502–508. 10.1038/s41586-022-04434-5

Tsekou, H., Angelopoulos, E., Paparrigopoulos, T., Golemati, S., Soldatos, C.R., Papadimitriou, G.N., Ktonas, P.Y., 2015. Sleep EEG and Spindle Characteristics After Combination Treatment With Clozapine in Drug-Resistant Schizophrenia: A Pilot Study. Journal of Clinical Neurophysiology 32, 159. 10.1097/WNP.0000000000000145

Tsuang, M.T., Faraone, S.V., 1995. The case for heterogeneity in the etiology of schizophrenia. Schizophrenia Research 17, 161–175. 10.1016/0920-9964(95)00057-S

van Os, J., Kenis, G., Rutten, B.P.F., 2010. The environment and schizophrenia. Nature 468, 203–212. 10.1038/nature09563

Walker, M.P., Brakefield, T., Morgan, A., Hobson, J.A., Stickgold, R., 2002. Practice with Sleep Makes Perfect: Sleep-Dependent Motor Skill Learning. Neuron 35, 205–211. 10.1016/S0896-6273(02)00746-8

Walker, M.P., Stickgold, R., 2004. Sleep-dependent learning and memory consolidation. Neuron 44, 121–133. 10.1016/j.neuron.2004.08.031

Wallwork, R.S., Fortgang, R., Hashimoto, R., Weinberger, D.R., Dickinson, D., 2012. Searching for a consensus five-factor model of the Positive and Negative Syndrome Scale for schizophrenia. Schizophr Res 137, 246–250. 10.1016/j.schres.2012.01.031

Wamsley, E.J., Shinn, A.K., Tucker, M.A., Ono, K.E., McKinley, S.K., Ely, A.V., Goff, D.C., Stickgold, R., Manoach, D.S., 2013. The Effects of Eszopiclone on Sleep Spindles and Memory Consolidation in Schizophrenia: A Randomized Placebo-Controlled Trial. Sleep 36, 1369–1376. 10.5665/sleep.2968

Wamsley, E.J., Tucker, M.A., Shinn, A.K., Ono, K.E., McKinley, S.K., Ely, A.V., Goff, D.C., Stickgold, R., Manoach, D.S., 2012. Reduced Sleep Spindles and Spindle Coherence in Schizophrenia: Mechanisms of Impaired Memory Consolidation? Biological Psychiatry, Functional Consequences of Altered Cortical Development in Schizophrenia 71, 154–161. 10.1016/j.biopsych.2011.08.008

Winkler, A.M., Ridgway, G.R., Webster, M.A., Smith, S.M., Nichols, T.E., 2014. Permutation inference for the general linear model. Neuroimage 92, 381–397. 10.1016/j.neuroimage.2014.01.060

