## Supplementary material for "A spectrum of altered non-rapid eye movement sleep in schizophrenia": Figure S1

Table S1 Group differences between SCZ and CTR in the full sample (Wave 1 and Wave 2 combined)

| EEG variables | Significant Cluster N | Cluster description |
|---|---|---|
| SS density | 1 | 33 chs, p=0.002 max t=-6 at P7 |
| FS density | 1 | 57 chs, p=3e-04 max t=-9 at C2 |
| SS amplitude | 1 | 53 chs, p=3e-04 max t=-6 at P6 |
| FS amplitude | 1 | 51 chs, p=3e-04 max t=-6 at C2 |
| SS ISA | 1 | 57 chs, p=3e-04 max t=-8 at C2 |
| FS ISA | 1 | 57 chs, p=3e-04 max t=-7 at F1 |
| SS duration | 1<br>2 | 14 chs, p=0.05 max t=4 at Fp1<br>21 chs, p=0.02 max t=-4 at P1 |
| FS duration | 1 | 57 chs, p=3e-04 max t=-7 at CZ |
| SS chirp | - | - |
| FS chirp | - | 31 chs, p=0.002 max t=-6 at POz |
| SS frequency | 1 | 24 chs, p=0.01 max t=-5 at FZ |
| FS frequency | - | - |
| SS coupl mag | - | - |
| FS coupl mag | 1 | 32 chs, p=0.002 max t=6 at PZ |
| SS coupl overlap | 1 | 57 chs, p=3e-04 max t=-8 at P3 |
| FS coupl overlap | 1 | 24 chs, p=0.006 max t=5 at PO4 |
| SS coupl angle | 1 | 35 chs, p=3e-04 max t=-7 at P7 |
| FS coupl angle | 1 | 22 chs, p=0.01 max t=-5 at TP8 |
| SS ph-fr coupl | 1 | 49 chs, p=3e-04 max t=-5 at FC6 |
| FS ph-fr coupl | 1 | 8 chs, p=0.05 max t=3 at AFZ |
| SO density | 1 | 56 chs, p=3e-04 max t=9 at O2 |
| SO duration | 1 | 54 chs, p=3e-04 max t=11 at CZ |
| SO slope | 1 | 55 chs, p=3e-04 max t=-9 at CZ |
| SO neg.peak amplitude | 1 | 15 chs, p=0.04 max t=-3 at FCZ |
| SO p-to-p amplitude | 1 | 14 chs, p=0.05 max t=-3 at FCZ |
| PSD | 1<br>2 | 720 chs, p=0.01 max t=-8 at C2 at 3Hz (1.5-5.75Hz)<br>848 chs, p=0.01 max t=-7 at C4 at 13.75Hz (9.5-15.75Hz) |
| PSI | 1<br>2 | 210 chs, p=3e-04 max t=-7 at AF4 at 11Hz (3-14Hz)<br>227 chs, p=3e-04 max t=7 at P3 at 11Hz (6-19Hz) |

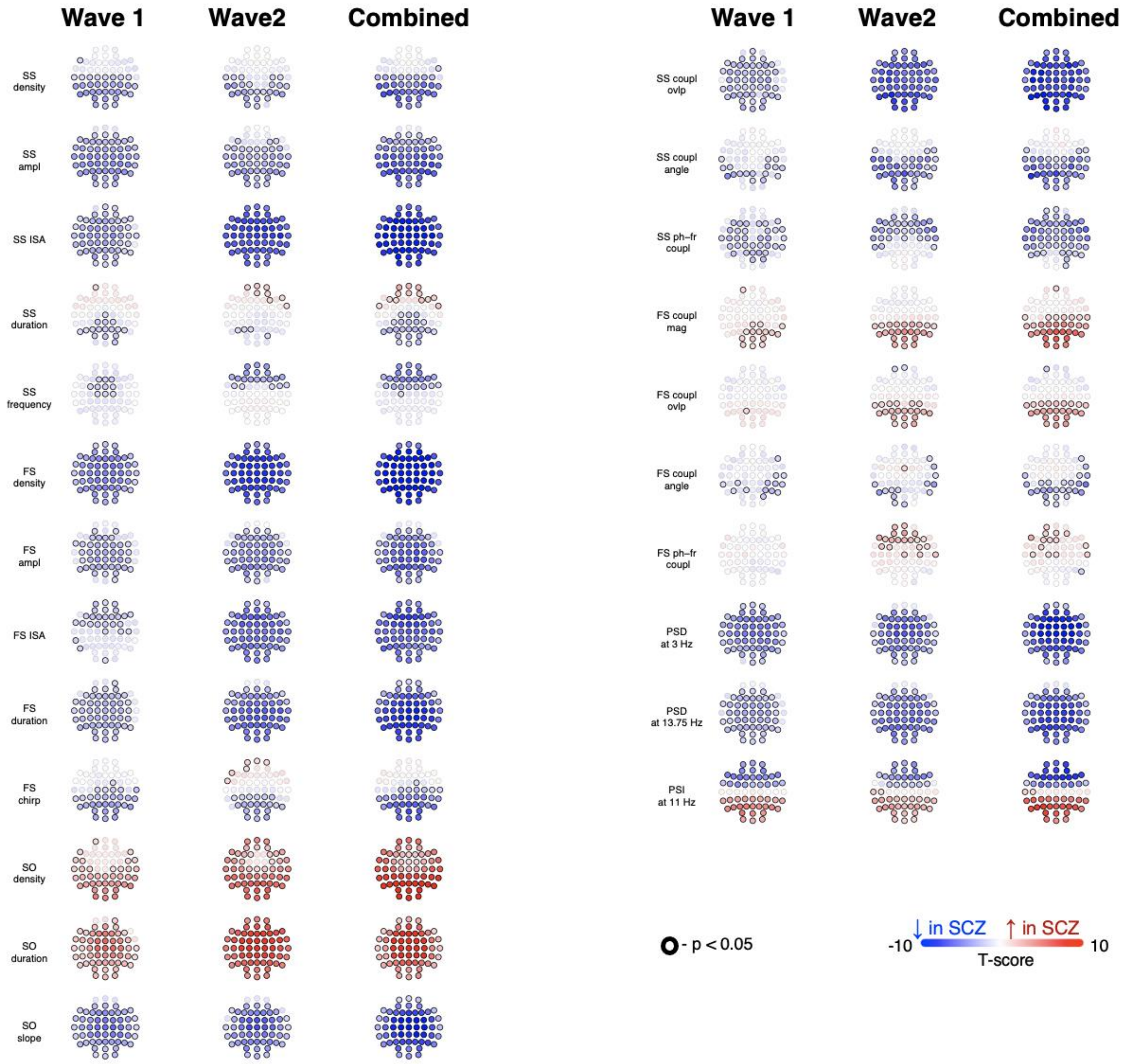


***Figure 1 Group differences between SCZ and CTR across waves and in the combined sample***

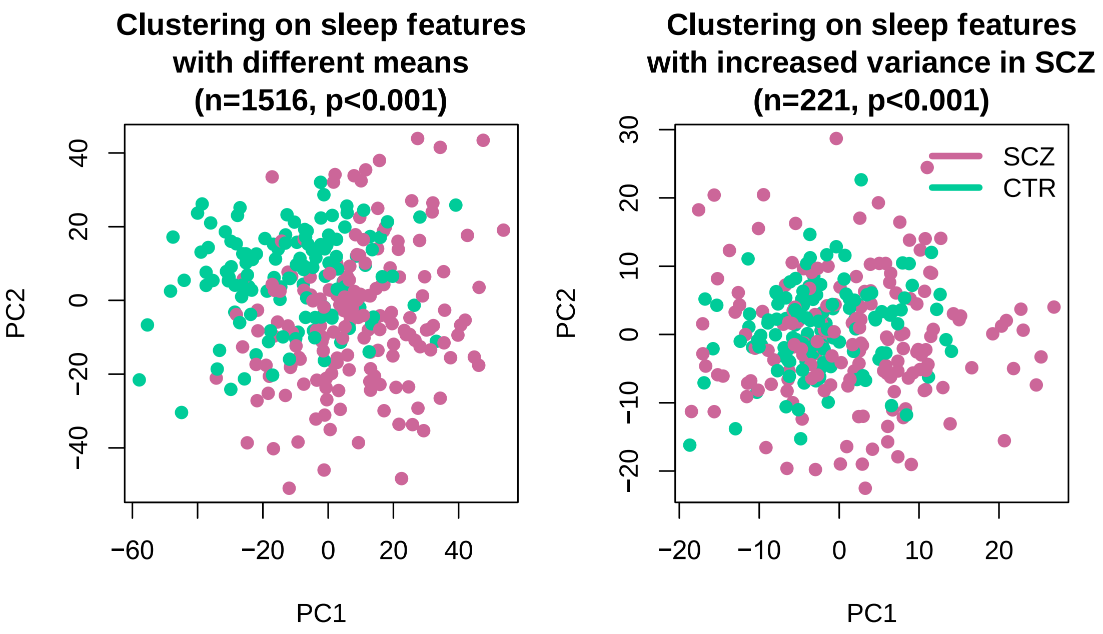


***Figure S2 No obvious sub-clusters within the SCZ group.*** *Visualization of the first two principal components as part of attempted clustering of individuals: the left plot – PCA based on group differences between means with p < 0.001 and the right plot – PCA based on all variables with ↑ variance in SCZ with p < 0.001*

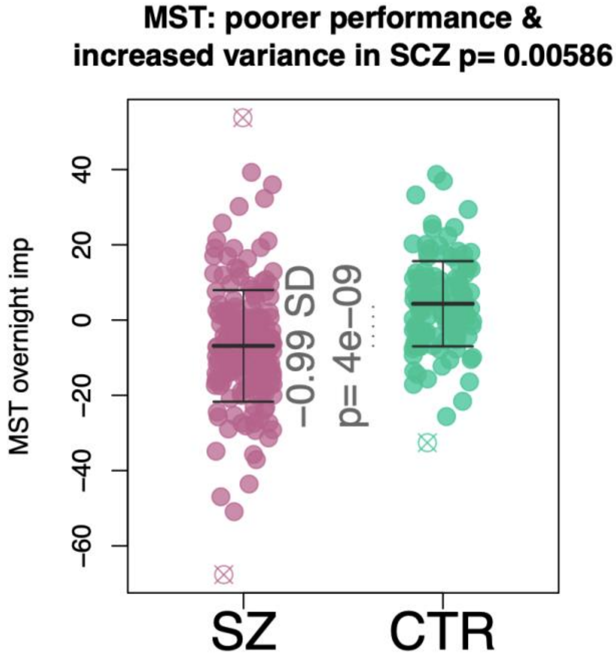


***Figure S3 Attenuation of overnight improvement of MST performance in the SCZ group versus controls.*** *Y axes represents the percentage increase in correct sequences from the best three training trials to the first three test trials the next morning*

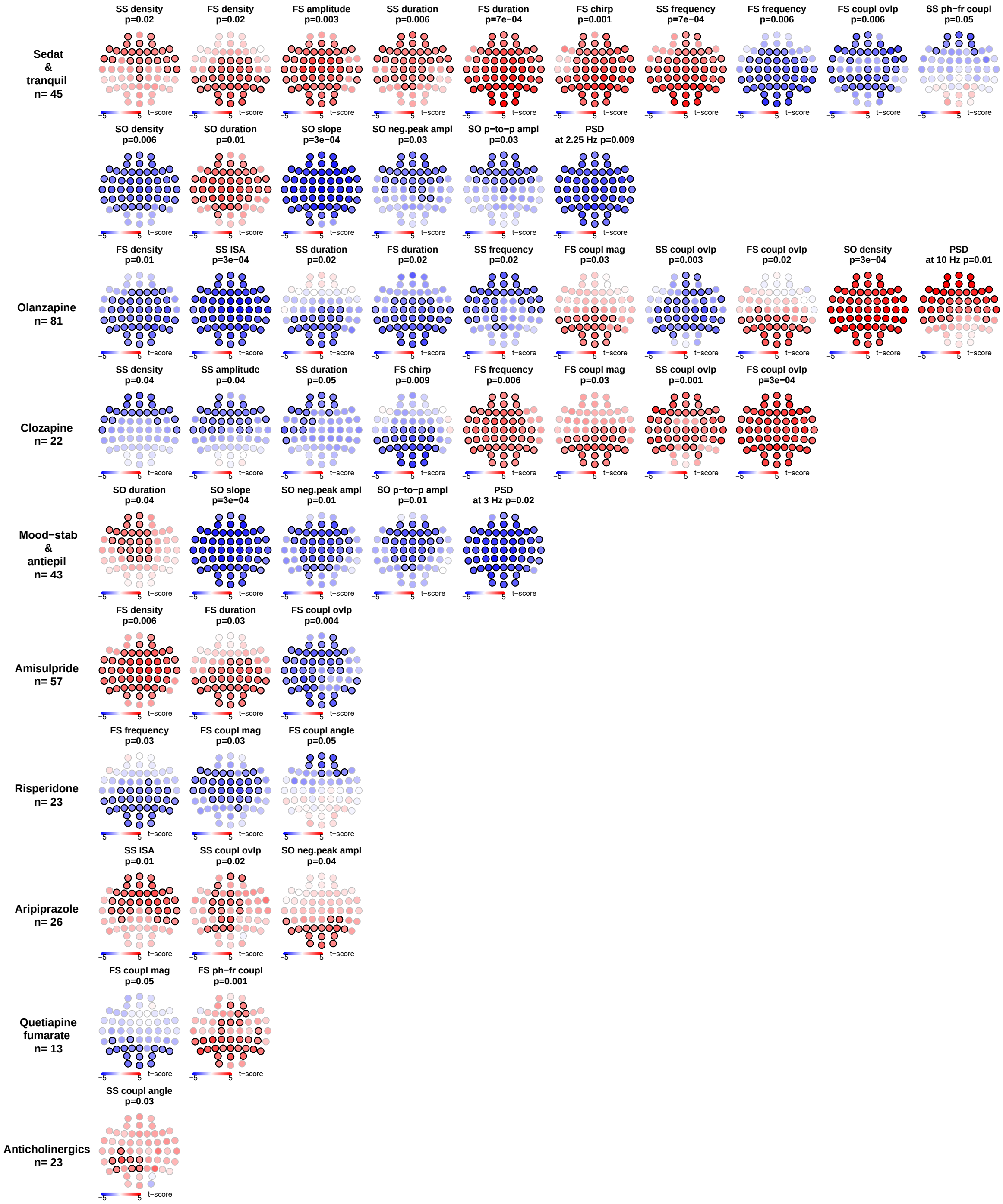


***Figure S4 Clusters of significant medication effects sorted by the number of significant clusters across sleep EEG variables***
